# Evolution and functioning of an X-A balance sex determination system in hops

**DOI:** 10.1101/2024.11.04.621975

**Authors:** Takashi Akagi, Tenta Segawa, Rika Uchida, Hiroyuki Tanaka, Kenta Shirasawa, Noriko Yamagishi, Hajime Yaegashi, Satoshi Natsume, Hiroki Takagi, Akira Abe, Miki Okuno, Atsushi Toyoda, Kyoko Sato, Yuka Honniden, Cheng Zhang, Koichiro Ushijima, Josef Patzak, Lucie Horáková, Václav Bačovský, Roman Hobza, Deborah Charlesworth, Takehiko Itoh, Eiichiro Ono

## Abstract

Chromosomal sex determining systems with male heterogamety include actively male-determining-Y and X-A balance systems, both of which are found in animals and plants. The sex-determining genes have been identified in several active-Y plant systems, but the evolution and functioning of X-A balance systems remains mysterious. To study this, we sequenced and compared the genomes of two hop species. The evolution of the hop X-A balance system involved an ancient recombination suppression event across a large X chromosome region shared by both species. In one species, an autosome fused to this ancestral sex chromosome, and recombination was subsequently suppressed again. The two evolutionary strata created in this neo-X have degenerated to different degrees, and evolved correspondingly different dosage compensation levels that correlate with histone modification patterns. Finally, we identified an X-specific ETR1-like ethylene receptor in the ancestral X region. Its dosage may affect sex determination, as part of the counting mechanism of this X-A balance system.

**One sentence summary:** Based on whole genome sequences of the cultivated hop, *Humulus lupulus*, and its wild relative *H. japonicus*, we describe the evolution of sex chromosomal regions, three of which that evolved region-specific dosage compensation, and identify a candidate gene involved in their X-A balance sex determining system.

## Introduction and background

In both animal and plants, independent lineages have evolved genetic sex determination systems and sex chromosomes. While changes between one genetic sex determining and sex chromosome system to another are documented in many animal lineages, recent changes from functional hermaphroditism to the state of having separate sexed individuals (dioecy) are more frequent in plants, making them suitable for studying how such systems evolve in the first place. Since the first finding of sex chromosomes in angiosperms in 1923, in *Humulus, Rumex*, and *Silene* (1-3), these have been studied using mainly genetic and cytogenetic approaches. These three genera all have highly heteromorphic sex chromosomes, allowing it to be determined that males are the heterogametic sex (4). The genus *Silene* has strongly dominant actively male-promoting, Y-linked factors, whereas *Humulus* and at least some *Rumex* species have systems in which the ratio of X chromosomes (X) and autosomes (A) determines sex, similar to the X-A balance systems in the model animal species, *Drosophila melanogaster* (5-6) and the nematode *Caenorhabditis elegans* (7). In such species, the X/A ratio determines sex, with a ratio of 0.5 or less determining maleness, 1 or more femaleness. The sex of genotypes with 0.5 < X/A < 1.0 differs between species; as demonstrated in experimentally produced polyploids with ratios between 0.67 and 0.75 in species of *Rumex* (8), and in *Cannabis* (hemp) and *Humulus* (hop), both in the family Cannabaceae (9-10), these are intersexes. The mechanisms are now understood in *D. melanogaster* and *C. elegans*, which have both evolved dosage compensation mechanisms that balance X-chromosome gene expression between the sexes, albeit with different systems for counting X chromosomes and different ways of adjusting expression (11).

Although Y-linked candidate sex determining genes have been identified in several plant species with active Y-systems (12-18), much less is known about plants with X-A balance systems. In some plant species with active Y-systems, two dominant Y-linked factors promote male functions, and one of them suppresses female functions (13, 15, 17, 19). Other systems involve a single Y-linked dominant male-promoting factor, either because variation at a second gene is no longer present (12), or because a turnover event has replaced an ancestral system with single gene control of sex, as is likely in the Salicaceae (16). The evolution of balance sex-determining systems has not been studied empirically, but sex is independent of the presence or absence of the Y-chromosome (reviewed by 4, 20). Data on the mechanisms of sex determination in different organisms are inadequate to reconstruct the history of these systems’ evolution (21). However, it is possible to test the prediction that balance systems should be older than those of species with active Y systems, even those with highly heteromorphic sex chromosomes suggesting that they evolved long ago. It is also of interest to investigate the counting mechanism by which the X/A ratio acts. This seems likely to evolve after profound degeneration and dosage compensation have evolved.

Cultivated hop (*Humulus lupulus*, a major crop used in brewing beer) is a member of a genus whose members are all dioecious and have X-A balance systems, as do species of *Cannabis*, in the same plant family, although the genera are distantly related (Fig. 1). *Cannabis sativa* and *H. lupulus* share the same karyotype (2n = 18 + XX/XY) and the same sex chromosome pair, and the sex determining system is thought to be shared (22). However, a wild relative, *H. japonicus*, whose genome size is about twice that of *H. lupulus* (23), has a sex chromosome-autosome fusion, resulting in the karyotype 2n = 14 + XX/XY_1_Y_2_ (Fig. 1a) (24). Here, we describe complete genome sequence assemblies of both these species, including their phased X and Y chromosomes, and initiate understanding of the X-chromosome counting mechanism.

**Figure 1.**
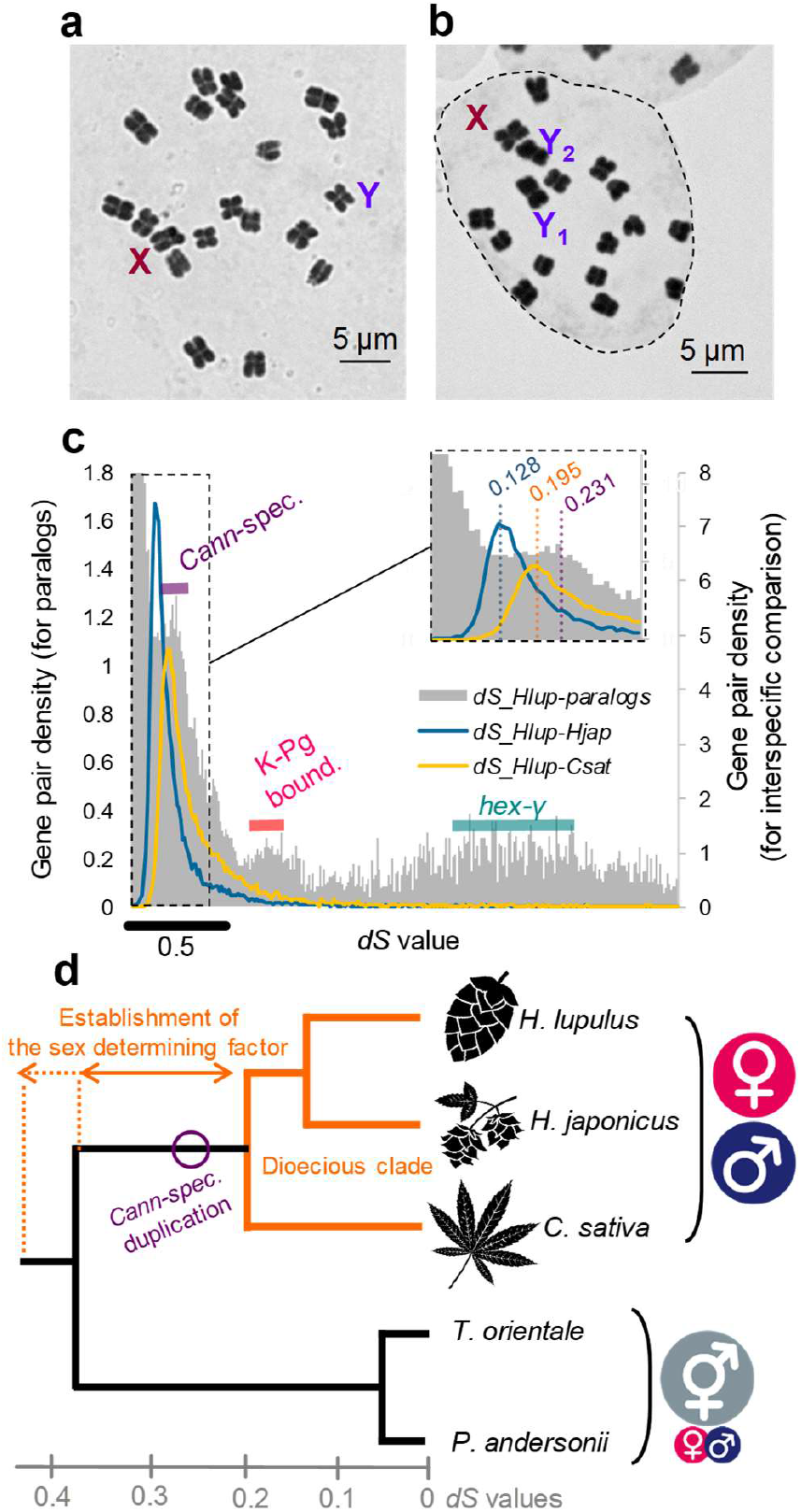
Sex chromosomes and genome evolution in the family Cannabaceae. **a-b**, Karyotypes of *H. lupulus* (**a**) and *H. japonicus* (**b**) males, showing the sex chromosomes. **c**, Distributions of *dS* values between paralogs in the *H. lupulus* genome, and of the interspecific (*H. lupulus-H. japonicus*, and *H. lupulus*-*C. sativa*) *dS* values using genes in the *H. lupulus* genome with complete coding sequences. The results suggest three whole genome duplication events, the most recent pre-dating the divergence of the genera *Humulus* and *Cannabis*. Cann-spec.: Cannabaceae-specific duplication, K-Pg bound.: plant-specific frequent genome-wide duplication approx. 66 mya with the last mass extinction, hex-γ: hexaploidization shared amongst most of the dicots plants. **d**, Schematic phylogeny showing the evolution of dioecy in the lineage that includes *Humulus* and *Cannabis*. The X axis shows *dS* values, which is roughly proportional to numbers of generations. The outgroup species *Trema orientale* and *Parasponia andersonii* are also members of the Cannabaceae, and are polygamous (mostly hermaphrodite), although we cannot exclude the possibility that they share the *Humulus* and *Cannabis* sex determining system. Few details of the sexual systems of members of the other 5 family Cannabaceae genera are known, but *Chaetachme* and *Pteroceltis* are described as dioecious. To be conservative, we label the the lineage that includes *Humulus* and *Cannabis* the “dioecious clade”, but that dioecy could have been established muvh earlier in the family.

### Genome sequences of *Humulus* species and genetic and physical maps

Chromosome-level whole genome sequences of a *H. lupulus* cv. Saaz female and a male from a cultivar named “10-12”, and a male *H. japonicus* from the “Seta-line” were determined using PacBio HiFi and Illumina Hi-C reads prepared with the Omni-C library kit. For *H. lupulus*, both pseudo-haploid and haplotype-resolved assemblies were constructed. The pseudo-haploid assemblies include sequences corresponding to 10 chromosomes (9 + X) for the Saaz female and 11 (9 + XY) for the 10-12 male, consistent with the karyotypes in this species. For the *H. japonicus* male, the pseudo-haploid assembly includes the expected 10 chromosomes (7 + XY_1_Y_2_). BUSCO evaluations indicated high completeness of conserved genes (> 98%) for all assemblies, and several chromosome sequences include both telomeres (tables S1-S4, fig. S1).

The distribution of silent site divergence (*dS*) between likely paralogs within *Humulus* peaks nears 23% indicating a whole genome duplication before *Humulus* and *C. sativa* diverged from their common dioecious ancestor (*dS* between orthologs in these genera show a peak just below 20%; Fig. 1b). The two *Humulus* species diverged considerably more recently, as pairwise *dS* values for orthologs are mostly around 12.8% (Fig. 1c). Fig. 1d summarizes the phylogeny and estimated times of these events.

All chromosomes in the *Humulus* genome assemblies (fig. S1) are larger than 100 Mb, and genetic mapping detected recombination only at the chromosome ends (or subtelomeric regions; ig. S2-3), consistent with previous information about recombination patterns in physically large chromosomes (reviewed by 25). As expected in recombinationally inactive regions, transposable element abundances tend to be high across much of the assemblies (figs. S4-5), and DNA methylation shows the inverse pattern (fig. S6-7). In *H. lupulus*, the left-hand approx. 35Mb region of the sex chromosome assembly recombines between the X and Y chromosomes in our F_1_ mapping population (fig. S2), consistent with this being a pseudo-autosomal region (PAR). In contrast, in *H. japonicus*, neither the Y_1_ nor the Y_2_ recombined with the X chromosome (see the legend of fig. S3), and linkage disequilibrium (LD) in a sample from a natural population in central Japan (see Methods for the details) is strong throughout both the Y_1_ and Y_2_ chromosomes, and between Y_1_ and Y_2_ variants (fig. S8); no PAR was detected in either the *H. japonicus* Y_1_ or Y_2_ assembly.

### Sex chromosomes of *Humulus* species

In the large pericentromeric fully Y-linked region of *H. lupulus*, Y-X synonymous site divergence (*dS*_*XY*_) estimates for coding sequences exceed 20%, suggesting that the system is slightly older than that in an active-Y system *Silene latifolia*, with a value of about 20% (17). A change-point test (26) identifies the region between 45 and 93 Mb as having been fully Y-linked since shortly before the split between *Humulus* and *Cannabis*, as *dS*_*XY*_ for the 19 genes identified in the region exceeds the divergence from *C. sativa* (*dS*_*HC*_ in Fig. 2a). This is therefore an ancient “Stratum 0”. The region distal to 93Mb is defined as a younger “Stratum 1”, as *dS*_*XY*_ values (145 genes) are similar to those for divergence from *C. sativa* (*dS*_*HC*_ in Fig. 2a). A PAR is defined based on (i) its terminal location and synteny (Fig. 2c) in the independently assembled terminal X and Y regions (before 45 Mb in the X assembly, and also with the independently assembled *H. japonicus* terminal X region, see Fig. 2f), and (ii) Y-X divergence rarely exceeding 10%, much lower than in the rest of the X (Fig. 2a). However, a 10-Mb part of this region is inverted between the Y and the X assemblies, and part of the left-hand end probably does not recombine, as Y-X synonymous site divergence estimates for its coding sequences are often around 5%, similar to the high values near the start of the large pericentromeric fully Y-linked region (fig. S9). Thus, recombination suppression may recently have evolved in hops, forming one or more evolutionary strata which we collectively term HlStratum 2 to distinguish this from the *H. japonicus* Stratum 2 described below. However, as a sex-linked region cannot evolve distal to a PAR, at least one inversion must have changed the arrangement after recombination stopped.

**Figure 2.**
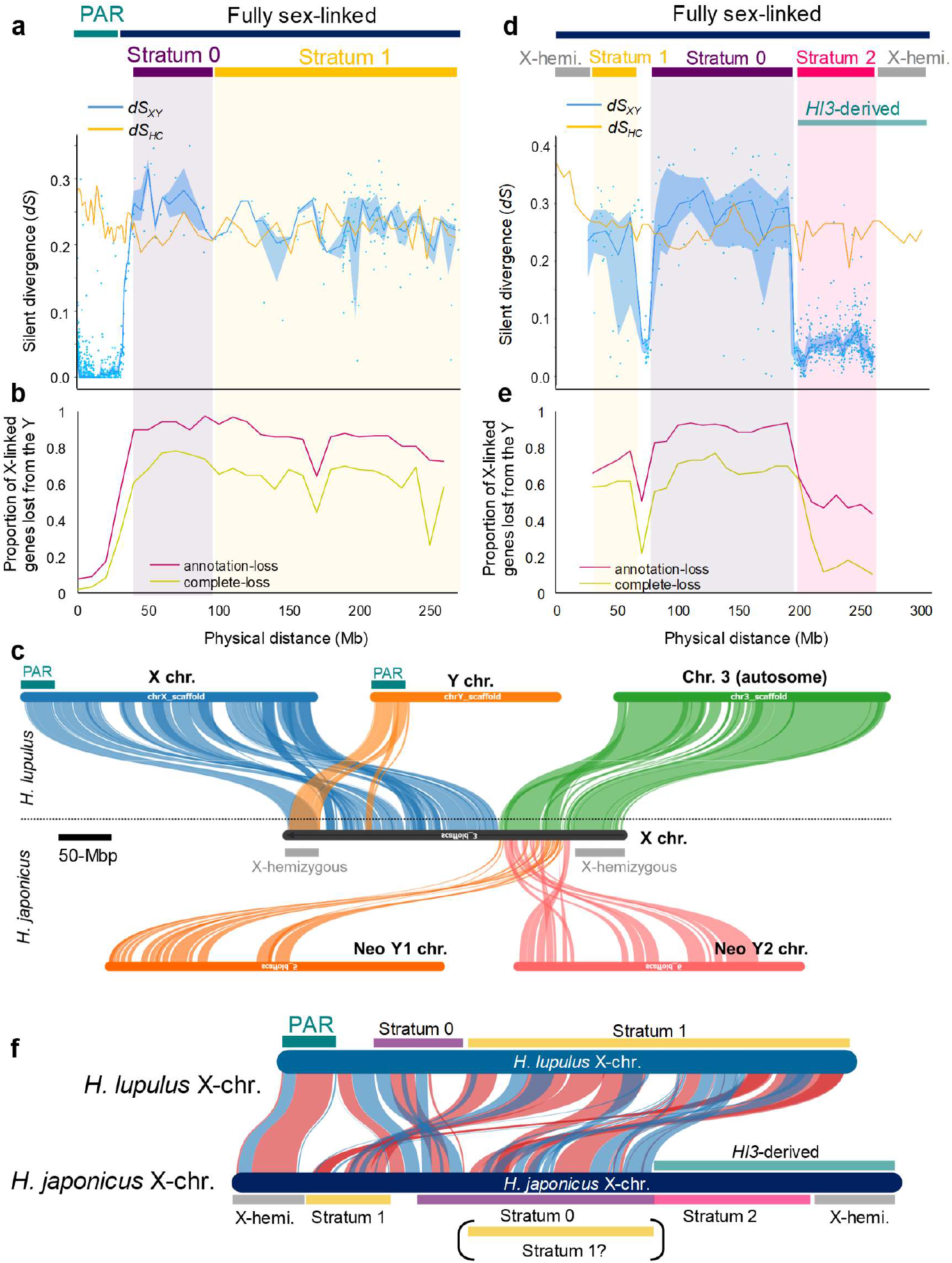
Structures of the sex chromosomes in the two *Humulus* species studied. **a**, Median values (with SD ribbon) of silent site per nucleotide divergence (*dS*) between *H. lupulus* X-and Y-linked alleles (*dS*_*XY*_) and between orthologs in *Humulus and Cannabis* (*dS*_*HC*_). The x axis shows the genes’ positions in the *H. lupulus* X chromosome physical map. The *dS*_*XY*_ values identify a large non-recombining region in which recombination stopped at about the same time as the divergence of the genera (confirmed by the results in the homologous *H. japonicus* region, see below). Because the left-hand part suggests a statistically earlier divergence time, we designate it Stratum 0, and the rest Stratum 1. The left-hand end of the *H. lupulus* X assembly forms a PAR. **b**, The proportions of X-linked genes lost from the *H. lupulus* Y, using two measures, annotation loss (defined as a gene for which no Y-linked copy was found by BLAST among the predicted genes) and complete loss (no similar sequence found by BLAST in the Y genome assembly). These proportions are similar across the Stratum 0-1 fully sex-linked region. **c**, Synteny based on the gene orders in the *H. lupulus* X/Y pair and chromosome 3 versus the *H. japonicus* X (derived from an X-autosome fusion involving chromosome 3) and the Y_1_ and Y_2_ chromosomes. The region corresponding to chromosome 3 shows fragments of synteny with both Y_1_ and Y_2_. **d**, Median *dS*_*XY*_ values in *H. japonicus*, again compared with *dS*_*HC*_ values. No PARs are detectable, as genes at both ends of the enlarged X chromosome retain no recognizable Y-linked alleles; their hemizygosity in males suggests that these regions are both highly degenerated, but their age cannot be estimated, due to the lack of XY gene pairs. In the remainder of the assembly, the ancient fully Y-linked region detected in the *H. lupulus* X is again evident, but the strata 0 and 1 detected by change-point analysis do not correspond to those in *H. lupulus* (see part **f** below). The rest corresponds to the ancestral chromosome 3, part of which forms a more recent stratum, 2, which is partially degenerated (part **e**), and part is entirely degenerated and hemizygous (as indicated in part **d), f**, Synteny between *H. lupulus* and *H. japonicus* X chromosomes (blue and red bands respectively indicate identical and inverted ordering). The X chromosomes of the two species are highly rearranged, with many inversions. The rearrangements are confined to the likely pericentromeric region, while the region corresponding to the *H. lupulus* PAR is syntenic, suggesting that this region continued to recombine in both species for a considerable portion of the time since they diverged.

Consistent with an ancient fully sex-linked region, in average about 70% of X-linked (presumably ancestrally present) genes have been completely lost from Y chromosome in the Stratum 0 region (Fig. 2b), and many more are incomplete (labelled “Annotation loss” in the figure”). Overall, nearly all genes in Strata 0 and 1 are either hemizygous or non-functional in males. An analysis of the gene orders (fig. S10) detects weak synteny with the Y chromosome across the first 95Mb of the X chromosome assembly, other than the PAR.

The *H. japonicus* sex chromosomes differ from those of *H. lupulus* by an X-autosome fusion involving chromosome 3, creating an enlarged X and two neo-Y chromosomes, Y_1_ and Y_2_, respectively corresponding to the *H. lupulus* X (*Hl*X), and chromosome 3 (*Hl*3; Fig. 2c). Values of *dS*_*XY*_ exceed 0.2 in *Hl*X, similar to those in *H. lupulus*, as expected from their shared ancestry (Fig. 1d). Part of the *Hl*3-derived region, “Stratum 2” in Fig. 2d, retains copies of most genes, and most neo-Y-X synonymous site divergence values are < 0.1, consistent with the fusion having occurred after the hop species split, long after recombination between the ancestral XY pair was suppressed. Within *Hl*X, change-point analysis detects significantly different *dS*_*XY*_ values, again defining two strata, but arranged in the opposite order from those in *H. lupulus* (Fig. 2f). Both strata (S0, from 80 to 190Mb in the *H. japonicus* assembly, and a younger S1 between 30 and 70Mb), retain a few Y-linked coding sequences, but have lost most Y genes (Fig. 2e). The region between 70 and 80Mb has much lower *dS*_*XY*_ values, and could reflect a translocated PAR.

Recombination between X and Y clearly stopped first in the ancestor of the *Hl*X and corresponding *H. japonicus* region, followed by the fusion with *Hl*3 in the *H. japonicus* lineage. Since the *H. lupulus* chromosome 3 has two terminal recombinationally active non-pericentromeric regions, the event presumably fused one such region to the non-PAR end of the ancestral X (Fig. 2f). Importantly, Y-X divergence across the Stratum 2 region adjoining the fusion breakpoint, between 200 and 270 Mb, indicates a complete lack of recombination, reflecting an additional recombination suppression event after or coinciding with the fusion. Interestingly, although Stratum 2 is considerably younger than the ancestral X strata, it is also degenerated, showing about 50% annotation loss and almost 20% complete loss (Fig. 2e). Degeneration within a time corresponding to synonymous site divergence of < 0.1 is surprisingly fast, given that purifying selection in the haploid pollen should hinder loss of function of Y-linked genes (27).

Within Stratum 2 of *H. japonicus, dS*_*XY*_ values are discontinuous, partially corresponding to Y_1_ and Y_2_ allelic blocks (fig. S11), suggesting that multiple rearrangements caused X-Y recombination to stop at various times; however, it is difficult to infer the detailed history, as rearrangements are also likely after recombination stopped. Of the total of approximately 600 Y_1_ and Y_2_ genes, most genes in the *Hl*3 Stratum 2 region are found in the rearranged Y_2_, which is therefore not derived by duplication of the Y_1_ chromosome. However, though some *Hl*3 genes are detected in Y_1_ (Fig. 2c). A few remaining Y gametologs in *Hl*X are found in Y_1_, but many more are in Y_2_, with four in both (figs. S11, S12). Most likely, after the fusion that created Y_2_, some regions or genes were duplicated from Y_1_. Duplications might be favoured if Y-linked copies of important *Hl*3 genes are losing function, as this can act as a form of dosage compensation.

It is puzzling that the *H. japonicus* X chromosome has male-hemizygous regions at both ends, especially as the left-hand end is homologous with the *H. lupulus* X PAR (Fig. 2f). The male hemizygosity of the right-hand distal region of this X assembly is also puzzling (Fig. 2d), as it indicates greater degeneration than in Stratum 2, implying that it stopped recombining before Stratum 2 evolved, although, as already noted, complete Y-linkage between the centromere-proximal region and a terminal one precludes more recent recombination cessation of an intervening region.

### Identifying the oldest sex-linked regions in the two *Humulus* species

The oldest non-recombining stratum, occupying 48Mb in the *H. lupulus* X chromosome assembly, should include any gene(s) involved in sex determination in the ancestor of *Humulus* and *Cannabis*. As these species have X-autosome balance systems, their Xs should share any genes involved. The two *Humulus* Xs share some syntenic regions, but with many rearrangements, and some regions and genes are found only in one species’ X (Fig. 2f, fig. S13a-b). The autosomes are not greatly rearranged between the two *Humulus* species (fig. S13), even though all chromosomes include large repetitive pericentromeric regions (fig. S2-5). These results suggest that rearrangements are promoted by sex linkage, and affect the X as well as the Y, albeit to a smaller extent. Fig. S13b shows that, after the *Hl*3 region became sex-linked, the neo-X rearranged between the two hop species more than did the autosomes in a similar time period. The LTR-type *Gypsy* class transposable elements show lineage-specific bursts on the X-chromosomes (fig. S14), which may contribute to their enlargement and rearrangement, similar to the evolution of the large and highly rearranged *Silene latifolia* X chromosome (17). In the *H. japonicus* sex chromosomes, mature tandem repeats were not detected in regions of CENH3 peaks detected by ChIP-seq analysis with a CENH3 antibody (Fig. S15), although, in another population, *Ty3/Gypsy* centromeric retrotransposons family strongly overlapped CENH3 signals (28). This suggests pericentric inversions within the rarely recombining X chromosome pericentromeric region of *H. japonicus*, changing the centromere positions between different populations.

### Dosage compensation in *Humulus*

When sex-linked regions degenerate, dosage compensation is expected to evolve. Partial compensation has been claimed in some plant sex chromosomes (29), though mechanisms are unknown. Transcriptomic data from young leaves in male and female *H. japonicus* siblings yielded clear evidence for complete dosage compensation in somatic tissues across both the large neo-X regions carrying male-hemizygous genes in (Fig. 3a). Stratum 2 showed less degeneration (with most genes still detectable on the neo-Y) and only a minority of genes show compensation (Fig. 3a), while the older Strata 0 and 1 (derived from the ancestral X region) are highly degenerated, and often show partial or complete compensation. These regions resemble the fully Y-linked regions, or MSY, in *H. lupulus*, which show similar Y degeneration levels throughout (Fig. 2a, Fig. S18).

**Figure 3.**
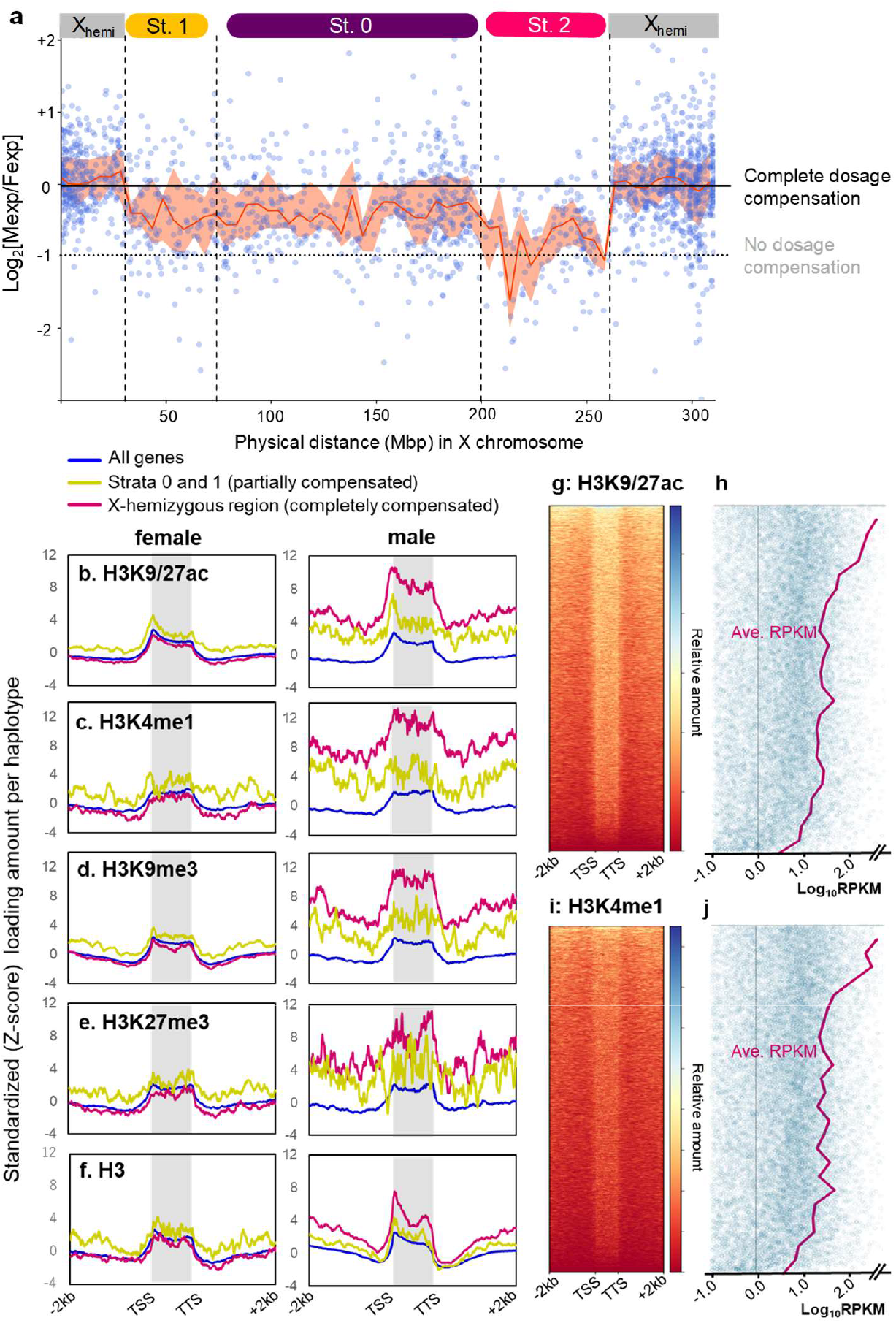
Regional dosage compensation in *H. japonicus* X chromosome. **a**, Distribution of expression bias in young leaves of male and female *H. japonicus* plants, shown as Log_2_(Male-expression/Female-expression), of genes, with their positions in the X chromosome physical map. Blue dots indicate expression biases in individual X-linked genes. Orange lines indicate median values in 5 Mb windows, with SD ribbons. **b-f**, Histone modification patterns in regions surrounding *H. japonicus*,genes, usinf three criteria, (i) whole gene marked (not compensated), (ii) Strata 0-1 (partially compensated), and (iii) X-hemizygous regions at both X-chromosome ends (completely compensated). TSS: transcription start site, TTS: transcription termination site. The typical euchromatic marks, H3K9/27ac (**b**) and H3K4me1 (**c**) are shown, and also H3K9me3 (**d**) and H3K27me3 (**e**), which often repress gene expression, exhibited statistically significantly higher loading in males than females in the X-hemizygous regions and Strata 0-1. The loadings correlate with the total H3 amount (**f**) in the X-hemizygous regions, whereas the H3 amounts in Strata 0-1 do not differ significantly between the sexes. **g-j**, Clustering of H3K9/27ac (**g**) and H3K4me1 (**h**) loading amounts in whole *H. japonicus* genes, which correlate positively with transcript levels (**i** and **j** corresponds to **g** and **h**, respectively).

The levels of dosage compensation do not correlate closely with DNA methylation status (fig. S16), but histone modification in and near genes (H3K4me1, H3K9/27ac, H3K9me3, and H3K27me3, and also total H3 amount) differ between the sexes and mirror the compensation levels, with especially high levels in males across both hemizygous compensated regions (Fig. 3b-f). In the partially compensated regions, total H3 amounts exhibited no substantial sex differences (Fig. 3f). H3K4me1 and H3K9/27ac are thought to mark euchromatin, and genome-wide analysis of their loading shows that they correlates positively with expression levels in leaves (Fig. 3g-j). These observations are consistent with male-specific upregulation of X-linked gene expression in the compensated regions, as in *Drosophila* species including the neo-X of *D. miranda* (30), especially those in which X-linked genes are hemizygous. In contrast, the other two epigenetic marks (H3K27me3 and H3K9me3, which often act as heterochromatic marks, and are also high in males in all the compensated regions) show no overall correlation with gene expression (fig. S17), and are unlikely to be related to dosage compensation.

### A candidate sex-determining gene

If *C. sativa* and *Humulus* indeed share X-linked gene(s) involved in X-autosome balance sex-determination, it may be possible to discover these by comparing the genome sequences, especially as (as mentioned above) the X chromosomes of *H. lupulus* and *H. japonicus* are highly rearranged, and the X chromosome arrangement of *C. sativa* is also different (Fig. 4a); overall, very few genes are in the oldest *Humulus* and *Cannabis* strata (which include parts of the two *Humulus* species’ Strata 0, and the right arm of the *C. sativa* (22); *N* = 51, Fig. 4b, table S5). Only one of these, an ETR1-like ethylene receptor gene, named *ETR1-like X-encoded Ethylene Receptor* (*EXER*), is found in all these species’ X chromosomes (fig. S19, S20, text S1). Ectopic expression of *H. japonicus EXER* under regulation by its native promoter (p*HjEXER-HjEXER*) resulted in feminization in *Nicotiana tabacum*, with some pleiotropic effects potentially explaining sexually dimorphic traits, including flower numbers (Fig. 4c-j, fig. S21, table S6). The pistil was elongated and the anthers were defective, producing mostly infertile pollen grains. Virus-induced gene silencing of *EXER* in female *H. japonicus* with apple latent spherical virus (ALSV) resulted in formation of anther-like structures in pistillate inflorescences and male-like inflorescence primordia (Fig. 4k-p). Supporting these results, treatment of female plants with silver thiosulfate (STS), an inhibitor of ethylene receptors, induced formation of functional male flowers (Fig. 4q, text S2, table S7-8, fig. S22). Furthermore, treatment with ethephon, which is metabolized to produce ethylene in plants, or ACC, a precursor of ethylene, resulted in formation of functional pistils by male plants (Fig. 4r-s, table S9). These results all suggest that the dosage of the ethylene receptor, *EXER*, could be an X-linked factor involved in this X-A balance sex determining system. Ethylene is involved in determining the sex of flowers in monoecious Cucurbits (31) and in the family Moraceae (32), which are close relatives of the family Cannabaceae. Furthermore, the dosage of *ETR1* ethylene receptor alleles can affect flower sex ratios in the monoecious *Cucurbita pepo* (33). These results suggest that the X-linked *EXER* (or *ETR1* ortholog) could be involved in these plants X-A balance sex determining system.

**Figure 4.**
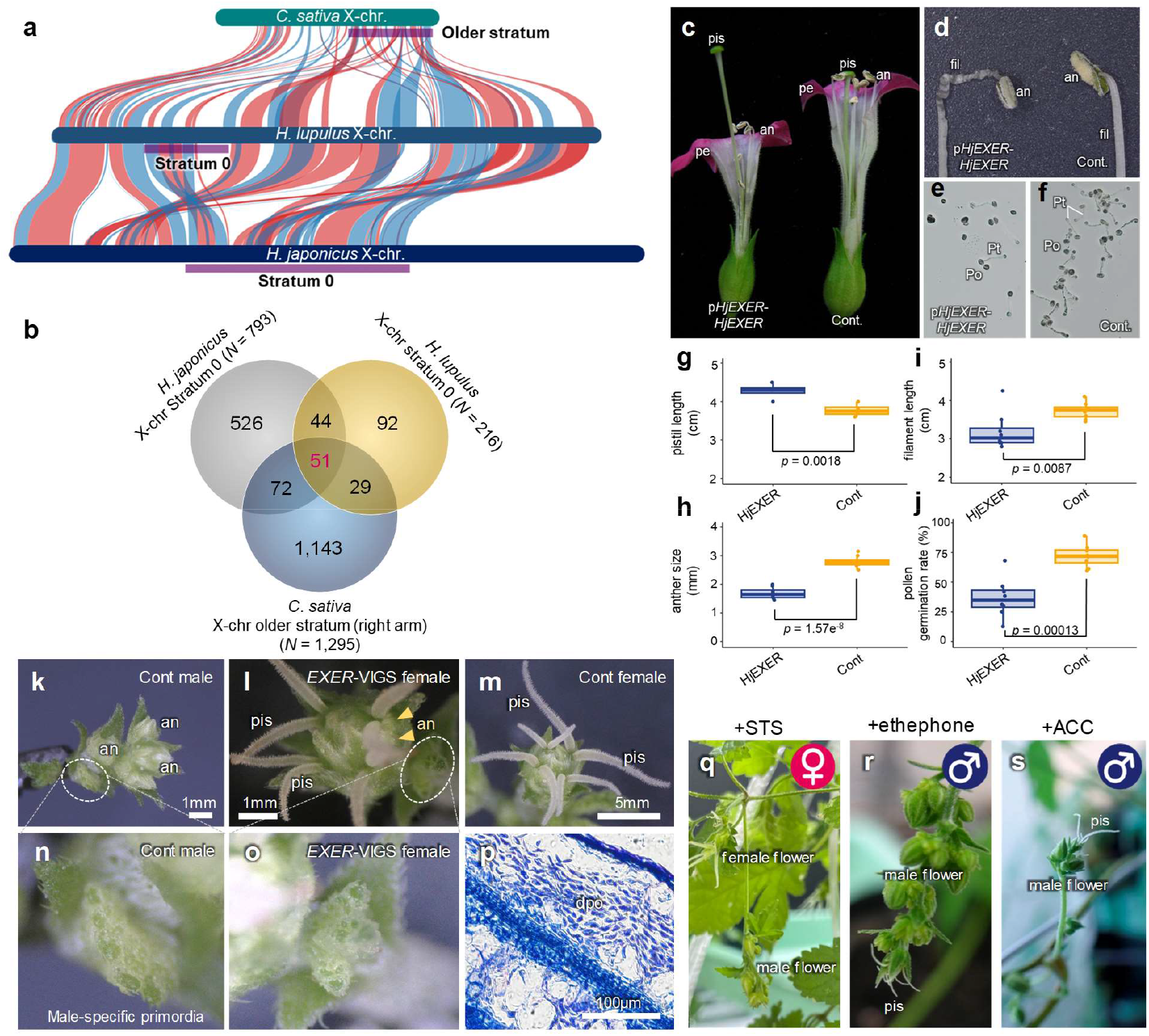
Identification of an X-linked sex determining candidate gene in the genus *Humulus*. **a**, Synteny of genes in X chromosomes of *C. sativa, H. lupulus*, and *H. japonicus*. Red and blue bands indicate syntenies with identical and inverted directions. The old strata (potentially including factors involved in sex determination are labeled with purple lines. **b**, Venn diagram of the genes included in the old strata in the three species. **c-j**, Results of transformation of *Nicotiana tabacum* with *H. japonicus EXER* (*HjEXER*) gene under the control of the native promoter (p*HjEXER-HjEXER*). The p*HjEXER-HjEXER* transgene caused feminization, with elongated pistils (**c** and **g**) and defective and shrunken anthers and filaments (**c, d, e-f, h-j**; an: anther, fil: filament, pe: petal, pis: pistil, po: pollen grain, pt: pollen tube), which produced less fertile pollen grains than control plants (**e-f, j**). **k-p**, Virus-induced gene silencing (VIGS) of *EXER* in female *H. japonicus*. **k**, Control male inflorescence. Inflorescences (**l**) of *EXER*-VIGS females produced anther-like structures (annotated by arrow heads) as well as pistils. **m**, Control female inflorescence. **n-o**, Control males and *EXER*-VIGS females often developed male flower (or inflorescence) primordia. **p**, Microscopic observation of dissected anther-like structures (panel **l** arrow heads) in an *EXER*-VIGS female. dpo: defective pollen grain. **q**, Effect of treatment with STS (an ethylene receptor inhibitor) in female *H. japonicus*, showing production of male flowers. **r-s**, Treatment with ethephon (an ethylene-inducer) (**r**), and ACC (an ethylene precursor) (**s**) in male *H. japonicus* induced female flowers with visible pistils.

### A model for evolution of the hop sex chromosomes

Our puzzling observations described above suggest the model in Fig. 5, for the evolution of the *H. japonicus* sex chromosomes. An ancestral degenerated Y chromosome evolved, carrying highly degenerated ancestral old strata that had evolved dosage compensation, and dosage-dependent sex-determination. The X subsequently fused with chromosome 3, creating a neo-X chromosome arm and a neo-Y_2_. Of the remaining Y-linked genes, most are present only in either the ancestral Y_1_ (with very few genes, some duplicated to/from Y_2_, see fig. S12) or in Y_2_, nearly all in Stratum 2; the hemizygous region and the younger Stratum 2 evolved after recombination stopped between the former chromosome 3 and Y_2_, perhaps involving closely linked polymorphisms controlling sexually dimorphic traits (reviewed in 10). Long recombinationally inactive regions in the middle parts of the ancestral chromosomes are probably the ancestral state, as the autosomes show this pattern (Fig. S2-3). Sexually antagonistic mutations in many genes would then be closely linked to the proto-Y-linked region, facilitating establishment of polymorphisms that would select for closer linkage. Once recombination stopped, TEs accumulated, expanding the non-recombining region. Our observations indicate that these processes occurred in both *Humulus* lineages with further gene losses after the X-autosome fusion, and evolution of dosage compensation in the new sex-linked region (fig. S14).

**Figure 5.**
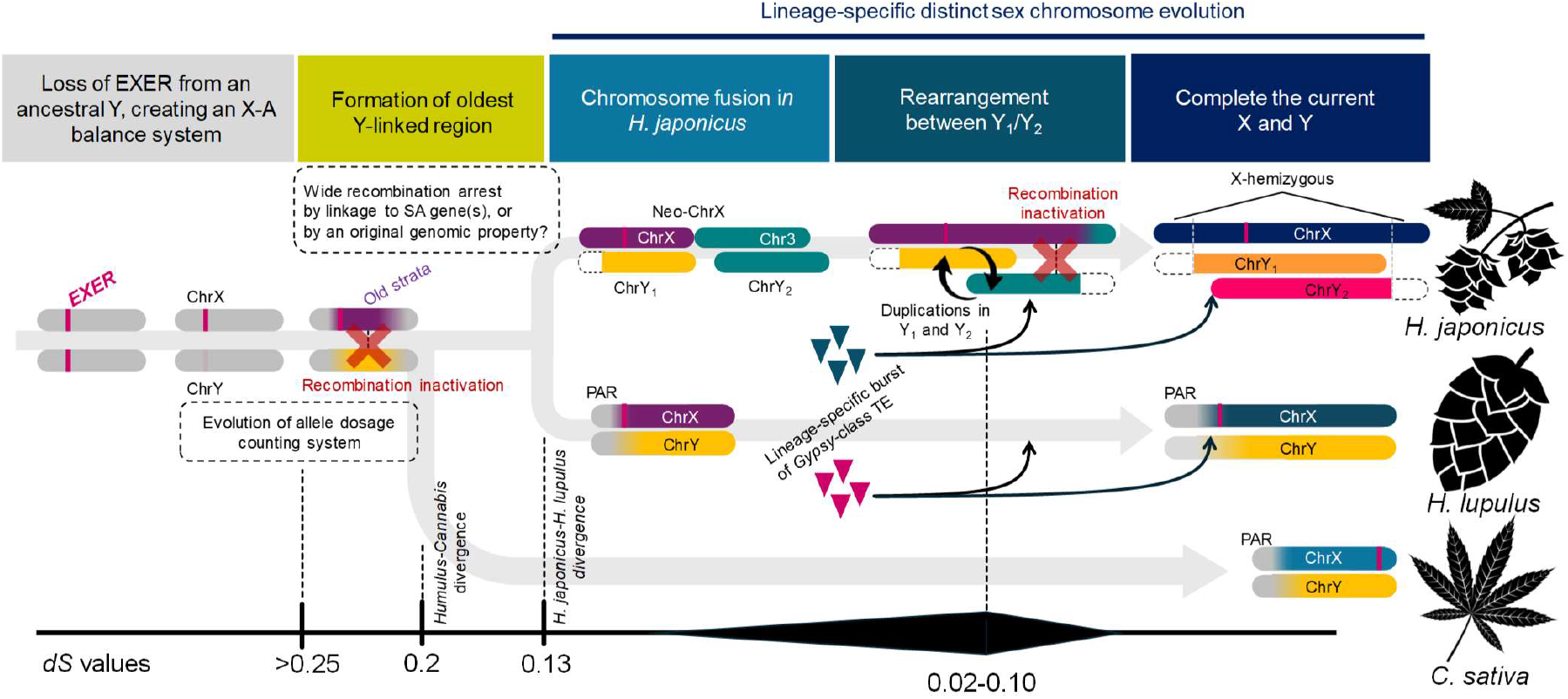
Model for the evolution of sex chromosomes in the genus *Humulus*.

## Acknowledgements

We thank Dr. Ryohei Terauchi (Kyoto University, Japan) and Drs. Makoto Suematsu, Fumihiko Sato, Hiroshi Kawashima, Hitoshi Matsubara, Hiroshi Shibata and all the members of Suntory Global Innovation Center Ltd. for helpful suggestions and dedicative support.

## Authors contributions

Conceptualization: TA, TI, EO

Methodology: TA, TS, RU, HT, KSh, NY, HY, SN, HT, AA, MO, AT, KSa, YH, CZ, KU, JP, LH, VB, RH

Investigation: TA, TS, RU, HT, KSh, NY, HY, SN, AA, MO, KSa, CZ, LH, VB

Visualization: TA, TS, HT, KSa, CZ, KU, TI

Funding acquisition: TA, KSh, RH, TI, EO

Project administration: TA, DC, TI, EO

Supervision: TA, DC, EO

Writing – original draft: TA, DC, TI, EO

Writing – review & editing: TA, DC, TI, EO

## Data and materials availability

All of the assembled genome sequences, the raw sequencing data and the annotated data were deposited to the DDBJ database (BioProject ID: PRJDB17941, PRJDB17942, and PRJDB18715). The sequencing data for reduced representation genome libraries, DNA methylome, ChIP-seq, and RNA-seq have been deposited in the DDBJ database (BioProject ID PRJDB19054, PRJDB19078, PRJDB19080, and PRJDB19097).

## Supplementary Materials

Materials and Methods Text S1 – S2

Fig S1 – S22

Table S1 – S10

References (33 – 70)

