## Supplementary Materials (methods, text S1-S2, figures S1-S22, and tables S1-S10) for "Evolution and functioning of an X-A balance sex determination system in hops": Supplemetary Materials (methods, texts, figures, and tables).pdf

**Title:**

**Running Title:**

Evolution of X-A balance sex determination in plants

**Authors:**

Takashi Akagi[1,2,*,**], Tenta Segawa[3*], Rika Uchida[1*], Hiroyuki Tanaka[4], Kenta Shirasawa[5], Noriko Yamagishi[6], Hajime Yaegashi[6], Satoshi Natsume[7], Hiroki Takagi[7], Akira Abe[7], Miki Okuno[4], Atsushi Toyoda[8], Kyoko Sato[9], Yuka Honniden[1], Cheng Zhang[1], Koichiro Ushijima[1], Josef Patzak[10], Lucie Horáková[11], Václav Bačovský[11], Roman Hobza[11], Deborah Charlesworth[12], Takehiko Itoh[4**], Eiichiro Ono[3**]
(*co-first authors, ** co-corresponding)

### Materials and Methods

#### Plant materials

Regarding *H. lupulus* (cultivated hops), the individuals for genome assembly, cvs. Saaz (female) and 10-12 (male) were grown in Suntory Global Innovation Center Ltd. Young leaves were harvested and immediately frozen with liquid nitrogen. Individuals for genetic mapping, F1 plants of *Humulus lupulus* were produced from a cross between the parents, Saazer (Osvald clone72/ a traditional cultivar, female) and 10-12 (male). F1 seeds were planted in the experimental field at Emma hop garden in Zatec, Czech Republic. For other experiments, including transcriptome analyses, the described F1 progeny was randomly selected to use experiments. Regarding *H. japonicas*, for genome assembly, we used an individual from the natural population, "Seta-line" harvested in the Seta region, Shiga, Japan (34.96° N, 135.91° E). This selection is maintained in Okayama University and Suntory Global Innovation Center Ltd. Young leaves were harvested and immediately frozen with liquid nitrogen. Individuals for genetic mapping, we used F1 population crossed within the Seta-line and also the Seta natural population. For dosage compensation analysis, including transcriptome and ChIP-seq analysis, we male and female siblings crossed within the Seta-line population.

#### Genome assembly and assessment

To improve and the quality of the *H. lupulus* draft genome previously published (34) and to conduct comparative genomic analysis in sex chromosomes, the HiFi reads for *H. lupulus* cv. Saaz (female), 10-12 (male) and a male *H. japonicus* were assembled with Hifiasm (35) in Hi-C mode to generate the initial contigs. In addition to the HiFi reads, Hi-C data was acquired for additional scaffolding purposes. The Hi-C library was prepared following the manufacturer's protocol using the Dovetail Omni-C Kit (Dovetail Genomics, Scotts Valley, CA, USA) and subsequently sequenced on a NovaSeq6000 instrument. The Hi-C scaffolding was performed using YaHS v1.2a.1 (36) with parameters "-q 1 –no-conitg-ec".

For the two *H. lupulus* plants, both pseudo-haploid and haplotype-resolved assemblies were constructed. For the haplotype-resolved assemblies (diploid genome) of *H. lupulus* cv. Saaz and 10-12, the input consisted of contigs constructed for each haplotype using Hifiasm (combined hap1.p_ctg and hap2.p_ctg), along with the results of mapping Hi-C data using the Dovetail Omni-C mapping pipeline (https://omni-c.readthedocs.io/en/latest/fastq_to_bam.html). Then, manual corrections were applied to the YaHS scaffolding results using Juicebox v1.11.08, leading to the final chromosome-level assemblies. The total lengths of the diploid genomes were 5.24 Gb with an N50 of 248.86 Mb for Saaz and 4.90 Gb with an N50 of 250.50 Mb for 10-12. The number of scaffolds was 3,074 for Saaz and 328 for 10-12, with the longest scaffolds being 309.43 Mb and 307.99 Mb, respectively. The diploid genomes included sequences corresponding to 18 autosomes and 2 sex chromosomes (X×2) for Saaz and 18 autosomes and 2 sex chromosomes (X, Y) for 10-12.

For the pseudo-haploid assemblies (haploid genome) of *H. lupulus* cv. Saaz, 10-12 and *H. japonicus*, the input consisted of contigs constructed for pseudo-haploid chromosome using Hifiasm (p_ctg), along with the results of mapping Hi-C data using the Dovetail Omni-C mapping pipeline. Then, manual corrections were applied to the YaHS scaffolding results using Juicebox v1.11.08, leading to the final chromosome-level assemblies. In addition, the two sex chromosomes (X, Y) of *H. lupulus* cv. 10-12 in the pseudo-haploid assembly were replaced with the corresponding sex chromosomes (X, Y) from the haplotype-resolved assembly. In the haploid genomes, the total lengths were and 2.52 Gb with an N50 of 247.17 Mb for Saaz, 2.67 Gb with an N50 of 250.77 Mb for 10-12, and 2.31Gb with an N50 of 245.10 Mb. The number of scaffolds was 2.67 Gb with an N50 of 250.77 Mb for 10-12, 164 for 10-12, and 379 for *H. japonicus*, with the longest scaffolds being 309.91 Mb, 309.74 Mb, and 313.10 Mb, respectively. The haploid genomes included sequences corresponding to 9 autosomes and 1 sex chromosome (X) for Saaz, 9 autosomes and 2 sex chromosomes (X, Y) for 10-12, and 7 autosomes and 3 sex chromosomes (X, $Y_1$, $Y_2$) for *H. japonicus*. The completeness of the assemblies was assessed using the BUSCO v5.4.7 (37). In BUSCO, the coverage rates of universal single-copy orthologous genes in eudicotyledonous plants (eudicots_odb10) were used as indexes of genome completeness. BUSCO analysis found 98.15%, 98.24%, 98.02%, 98.28%, and 98.20% of universal single-copy orthologous eudicotyledonous plant genes in the pseudo-haploid assemblies of Saaz (haploid), Saaz (diploid), 10-12 (haploid), 10-12 (diploid), and *H. japonicus* (haploid), respectively (table S1).

Telomeric repeats were identified using telomere identification toolkit (tidk) v0.2.41 (https://github.com/tolkit/telomeric-identifier). We searched "TTTAGG" sequences at the ends of the chromosomes. In the pseudo-haploid assemblies, telomeric repeats were found on 3 of the 10 chromosomes in Saaz (haploid), 1 of the 20 chromosomes in Saaz (diploid), 5 of the 11 chromosomes in 10-12 (haploid), 6 of the 20 chromosomes in 10-12 (diploid), and 1 of the 10 chromosomes in *H. japonicus* (haploid) (tables S2-4).

**TE and gene annotation**

A custom repeat library was created to identify repetitive sequences in the five *Humulus* assemblies. Specifically, de novo repeat libraries were constructed from whole-genome sequences using RepeatModeler v2.0.3 (38) and the LTR structure discovery pipeline. The five *Humulus* assemblies were then masked using RepeatMasker v4.1.2-p1 (39) with the custom repeat library.

Gene structural annotation of all *Humulus* assemblies was conducted using a combination of RNA-seq transcriptome sequences for gene structure prediction, homology-based gene prediction using protein sequences of related species and ab initio gene prediction. For the transcriptome-based gene prediction, RNA-seq reads were filtered by Platanus_trim (http://platanus.bio.titech.ac.jp/pltanus_trim) to remove adaptor and low-quality sequences. De novo transcriptome assembly was performed with Trinity v2.8.4 (40) and Oases v0.2.8 (41) and redundant sequences were trimmed with CD-HIT v4.6 (42). The assembled sequences were aligned using Gmap v2019-02-15 (43) and complete open reading frames (ORFs) were predicted by TransDecoder v5.0.2 (https://github.com/TransDecoder/TransDecoder). In addition to de novo assembly, gene prediction was also performed by mapping the RNA-seq reads to the genome. The filtered RNA-seq data were mapped to the genome sequences using HISAT2 v2.2.1 (44), and the mapped transcripts were assembled using StringTie v2.1.7 (45). ORF regions were then identified with

TransDecoder. For the homology-based method, protein sequences of *Cannabis sativa* (NCBI accession No: GCF_900626175.2), *Parasponia andersonii* (NCBI accession No: GCA_002914805.1), and *Trema orientale* (NCBI accession No: GCA_002914845.1) were used. They were aligned to the genome sequences using Spaln v2.3.3 (46) to predict gene structures. For the ab initio-based method, AUGUSTUS v3.3.2 (47) and SNAP v2006-07-28 (48) were used. Both programs were trained on gene models derived from 1,000 genes predicted by the transcriptome-based method. Finally, all predicted gene candidates were merged using the GINGER pipeline (49). Annotation results predicted 25,979 genes in Saaz (haploid), 52,194 genes in Saaz (diploid), 26,490 genes in 10-12 (haploid), 49,594 genes in 10-12 (diploid), and 26,320 genes in *H. japonicus* (haploid). The completeness of the predicted genes was evaluated using BUSCO v5.4.7 (37) with single-copy orthologs from the eudicots_odb10 database. BUSCO analysis found 98.54%, 98.84%, 98.75%, 98.88%, and 97.54% of universal single-copy orthologous eudicotyledonous plant genes in the gene models of Saaz (haploid), Saaz (diploid), 10-12 (haploid), 10-12 (diploid), and *H. japonicus* (haploid), respectively.

**Linkage maps in *H. lupulus* and *H. japonicus***

In *H. lupulus*, The 250 F1 plants were subjected to Genotyping by Random Amplicon Sequencing-Direct (GRAS-Di) analysis (50) and co-dominant markers were identified with GRAS-Di software v 1.0.5 (Toyota, Aichi, Japan). The GRAS-Di procedure was conducted by Genebay Co., Ltd. (Yokohama, Kanagawa, Japan).

To construct a genetic map, sequence data from 10-12 reference, excluding the Y chromosome, were aligned using minimap2 version 2.14 with F1 Grass di data from Saaz, 10-12, and 250 F1 progenies (51). The sequence alignment files were converted to BAM format using Samtools version 1.20, and SNP calling was performed using BCFtools version 1.13 (MQ >= 60, BQ >= 13) (52). The hop genetic maps were constructed using SNP markers, including 3,265 10-12-specific heterozygous polymorphisms.

SNP selection followed these criteria: (1) 10-12 specific heterozygous SNP loci, regions with a sequencing coverage depth >= 10 and heterozygous SNP allele frequencies between 0.3 and 0.7 were selected. (2) Loci with more than 5% missing data in the F1 population were excluded. (3) SNPs showing segregation consistent with a 1:1 ratio based on a Chi-square test of independence were selected. The selected SNPs were manually corrected to account for potential genotype errors, such as double recombination events, and to ensure accurate selection. These SNPs were then used to plot the physical and genetic distances.

To construct a genetic map in *H. japonicus*, the 96 F1 plants were subjected to MIG-seq analysis, according to the previous report (53). The sequenced reads were aligned to the *H. japonicus* reference genome using the same method as used for constructing the genetic map of *H. lupulus*, followed by SNP calling. The *H. japonicus* genetic maps were constructed using SNP markers, including 1,066 female parent-specific and 908 male parent-specific heterozygous polymorphisms. SNP selection followed a similar procedure as that used for hop, but for the $Y_1$ and $Y_2$ regions, SNPs were selected based on a sequencing depth of 0 in the female parent and a SNP allele frequency of 1 in the male parent.

For natural population of *H. japonicus*, the ddRAD-seq (54) data from 70 individuals of the natural population were aligned to the *H. japonicus* reference genome using the same method as for hop, followed by SNP calling. Positions with more than 5% missing data or a minor allele frequency (MAF) below 0.3 were excluded from the analysis. A total of 3,720 SNPs were used to generate the LD plot.

#### Synteny analyses

Gene-order-based synteny was detected with MCScanX (55), in which the detected collinearity was visualized using SynVisio (https://synvisio.github.io/#/). BLASTP analyses found homology, among the protein sequences in the X and Y chromosomes, or between two whole genomes (*H. japonicas* and *H. lupulus*), with an e-value cut-off of <1e$^{-20}$ for detection of the X-Y alleles, and <1e$^{-50}$ for the intragenomic comparison of *H. japonicas* and *H. lupulus*, with max_target_seqs = 2. Syntenic blocks were defined by using MCScanX, with BLASTP results and gff data. Sequence-based synteny analysis was performed using minimap2 and visualized with dotPlotly (https://github.com/tpoorten/dotPlotly), with a minimum alignment length of 10-50 kb.

#### Detection of genetic diversity

Silent divergence (*dS*) among the orthologous or paralogous gene pairs (<e$^{-20}$ in BLASTP analyses, max_target_seqs = 2), was aligned in-codon frames by using Pal2Nal and MAFFT ver. 7 under the L-INS-i model, according to the previous method (56). The in-codon-frame alignments were analyzed with MEGA X (57) to calculate the Jukes and Cantor corrected values of *dS*. To detect statistic change points in *dS* transitions, in the X chromosome of *H. japonicus* or *H. lupulus*, Pettitt's change-point test was conducted, with "pettitt.test" function in the "trend" package of R. Proportion of X-linked genes lost from the Y chromosome, which was defined as the rate of X-specific genes (or Y-lost genes), was evaluated with two criteria: (i) complete loss and (ii) annotation loss. Annotation loss is the situation that Y chromosome has no predicted protein-coding gene homologous to an X gene, in blastp analyses (<e$^{-20}$ for the threshold). Complete loss is the situation that the genomic sequences of Y chromosome have no homologous (<e$^{-20}$ for the threshold) counterpart to an X gene. In blastp analysis using an X gene as a query, if an autosomal gene (or sequences) preferentially hit rather than a Y gene (or sequences), we counted this as a gene lost from Y chromosome.

#### Detection of dosage compensation in X chromosome

Male and female young leaves (approx. 8-12$^{th}$ leaves from 5 biological replicates) in a sibling of *H. japonicus* or cvs. Saaz and 10-12 of *H. lupulus*, were harvested for transcriptome analyses. mRNA-seq libraries were prepared with a modified protocol of the previous study (17). Total RNA was isolated using PureLink Plant RNA Reagent (Invitrogen). mRNA was purified using the Dynabeads mRNA purification kit (Life Technologies). mRNA-seq library was constructed with KAPA RNA HyperPrep kit (Kapa Bioscience), followed by a DNA cleanup step with AMPure XP beads (Beckman Coulter; AMPure:reaction, 0.8:1). The constructed mRNA-seq libraries were sequenced on Illumina HiSeq X. All Illumina sequencing was conducted at Macrogen Japan. Sequencing reads were preprocessed using Python scripts (https://github.com/Comai-Lab/allprep/blob/master/allprep-13.py), to remove the adapter sequences and cut off the N or low-quality residues. The processed reads were mapped to the whole genes of *H. japonicus* or *H. lupulus*, using the Burrows-Wheeler aligner (BWA)-mem (http://bio-bwa.sourceforge.net/) with the default parameters. SAM files were generated from the mapping information and converted into read counts data, according to the previous report (11). To characterize dosage compensation in the X chromosome. we filtered genes in the X-hemizygous genes and X-genes lost from the Y chromosome in the criteria of annotation loss, as described above. Standardized expression levels (read numbers per kilobase and millions mapped reads: RPKM) were detected in each male and female group, and their biases were plotted in the X chromosome physical map.

**DNA methylome and ChIP-seq library**

With gDNA extracted from young leaves (8-12$^{th}$ leaves from 5 biological replicates for each male and female), DNA methylome libraries were prepared, according to previous reports (58-59). Briefly, gDNA was fragmented into *ca*. 250-350bp using Bioruptor-One sonicator (Sonic Bio) and end-repaired followed by adenosine-tailing steps. The fragments were ligated with a cytosine-methylated adapter. These procedures were conducted by using KAPA HyperPlus Kit (KAPA Biosystems). The adapter-ligated DNA fragments were purified with AMPure XP beads (Beckman Coulter), and a bisulfite conversion step was conducted with the EZ DNA Methylation-Gold kit (Zymo Research, USA). The bisulfite-converted DNA fragments were enriched by 8 cycles of PCR.

ChIP-seq analyses for assessment of histone modification was conducted according to the modified protocol of Akagi et al. 2024 (17). Total 0.5g leaf of a male or female *H. japonicas* was ground into fine powder with liquid nitrogen, and crosslinked with 1% formaldehyde for 10 min at room temperature. Crosslinking was quenched by the addition of glycine to a final concentration of 125 mM. Extraction of nuclei and immunoprecipitation was conducted according to Akagi et al. 2024. The chromatin precipitate was resuspended in 150 μL SDS lysis buffer (50 mM Tris-HCl pH 8.0, 10 mM EDTA, 1% SDS) and sheared using Bioruptor-One sonicator (Sonic Bio) for 8-16 cycles of ON/OFF for each 30 seconds at 10 °C. After centrifugation, 100 μL of the supernatant was used for immunoprecipitation. Immunoprecipitation was performed with 2-3 μg of anti-H3K9me3, -H3K4me1, -H3K9/27ac, -H3K27me3 antibodies (MA308A, MA302A, MA310A, MA323A; TaKaRa, Japan), or anti-H3 antibody (MA301A; TaKaRa). After washing, immune complexes were eluted with elution buffer (10 mM Tris–HCl, pH 8.0, 0.3 M NaCl, 5 mM EDTA, 0.5% SDS) and DNA was reverse-crosslinked by incubating overnight at 65 °C. DNA treated with proteinase K and RNase, was purified by phenol-chloroform method. Total 0.5-2.0 ng of purified DNA was applied to Illumina library preparation with the KAPA HyperPlus Kit (KAPA Biosystems).

The Illumina libraries were sequenced using Illumina's HiSeqX (Macrogen Japan). Raw Illumina reads were processed using custom Python scripts (http://comailab.genomecenter.ucdavis.edu/index.php/), according to the previous study (11).

Detection of methylated-cytosine in the DNA methylome data was performed with the methylpy pipeline (https://github.com/yupenghe/methylpy, (60). The averaged weighted methylation levels (detected separately for CG, CHG, and CHH) surrounding the gene were visualized with deeptools2 (61).

For ChIP-Seq reads were preprocessed and aligned to the *H. japonicas* genome sequences, with Burrows-Wheeler Aligner (BWA)-mem (62), with the default parameters. The standardized coverages surrounding the gene bodies were calculated and visualized with deeptools2. Furthermore, the standardized coverages surrounding the gene bodies were hierarchically clusterized with deeptools2 (using plotheatmap) to detect correlation to the expression levels in young leaves.

**Centromeric repeat assessment**

We mapped the ChIP-seq data with *H. japonicas* and *H. lupulus* CENH3 antibody (28) to the *H. japonicas* and *H. lupulus* whole genome sequences, respectively. Approximately 10Mb genomic regions surrounding putative CENH3-peak regions were extracted from the sex chromosomes. With StainedGlass package (63), we generated sequence identity heat maps to analyze tandem repeat distributions using a 2-kb window with the mm_f option set to 10 kb.

**Virus induced gene silencing with ALSV in *H. japonicus***

Binary plasmid vectors pCALSR1 and pCALSR2mL3mR3 (64-65), which encode RNA1 and RNA2 genomes of apple latent spherical virus (ALSV), respectively, were used for the generation of ALSV-EXER vector. A 201-bp fragment of the *H. japonicus* ETR1 ortholog, or *EXER* gene was amplified by PCR from genomic DNA of *H. japonicus* (Seta-line) with R2m3hpRecB-IFF and R2m3hpRecB-IFR (table S10) using PrimeSTAR GXL DNA Polymerase (TaKaRa Bio). To introduce the 201 nt fragment into the XSB cloning site of ALSV-RNA2 vector in-frame, pCALSR2mL3mR3 was linearized by inverse PCR with R2m3-ivF and R2m3-ivR (table S10) using PrimeSTAR GXL DNA Polymerase (TaKaRa Bio). The 201-bp insert fragment was introduced into the linearized vector using In-Fusion Snap Assembly Master Mix (Takara Bio) according to the manufacturer's instructions. The resulting plasmid, pCALSR2m3-EXER and pCALSR1 were independently introduced into *Agrobacterium tumefaciens* strain GV3101. Agroinoculation of *N. benthamiana* with the mixture of agrobacteria suspensions carrying pCALSR1, pCALSR2m3-hpREC, or pBIN:P19 (expressing tombusvirus silencing suppressor P19) was performed as described previously (64). The upper leaves of agroinoculated *N. benthamiana* were then used for rub-inoculation of *Chenopodium quinoa* to propagate ALSV-EXER vector.

Biolistic inoculation of *H. japonicus* with ALSV-EXER vector RNA was performed as described previously (66). In summary, total RNA was extracted from *C. quinoa* leaves infected with ALSV-EXER vector by TRI-Reagent (Sigma-Aldrich) and was coated onto 0.6 μm Gold Particles (Bio-Rad Laboratories, Hercules, CA, USA). The germinated *H. japonicus* cotyledons were bombarded eight times with the RNA-gold particles (5μg of RNA per shot) at the helium pressure of 20 psi. After bombardment, the plants were cured in a plastic dish at 18°C for one day under dark conditions and seven days under continuous light conditions. After curing, the plants were transferred to soil and grown at 25°C in a growth chamber under 16-h light/8-h dark conditions.

**Transformation of *Nicotiana tabacum***

Approx. 2kb promoter sequences and coding sequences (CDS) of the *H. japonicus* ETR1 ortholog, or *EXER*, were amplified from genomic DNA of a *H. japonicus* male individual (a Seta line maintained in Okayama University) by PCR using PrimeSTAR Max (TaKaRa Bio) (table S10 for primer information). The PCR amplicons of the *EXER* promoter region and CDS were separately cloned into the *HpaI* and *BamHI* sites of pPLV02 vector, respectively (67) to place the genes under the native promoter, to design *pHjEXER::HjEXER* construct. For cloning, we used the InFusion (TaKaRa Bio) pipeline, according to the manufacture's instruction. Tobacco plants (*Nicotiana tabacum*) cv. Petit Havana SR1 were grown *in vitro* under white light with 16-h-light and 8-h-dark cycles at 24°C. Young leaves of *N. tabacum* plants were transformed with *pHjEXER::HjEXER*-introduced *A. tumefaciens* strain EHA105, according to the method as previously described (13). Transformed plant tissues were selected with 100 μg/mL kanamycin on Murashige and Skoog (MS) medium. Rooted transgenic plants were transplanted into non-autoclaved soils, and grown in 16-h-light and 8-h-dark cycles at 26°C.

**Treatment of chemical compounds involving ethylene receptability and biosynthesis**

For female *H. japonicas*, young plants before flowering (planted in 100mm x 80mm pot) were sprayed with approx. 5mL of 2mM sodium thiosulfate (STS; 1% Chrysal K20-C diluted by distilled water) four times, once every week. For male *H. japonicas*, young plants before flowering (planted in 100mm x 80mm pot) were treated with 1mM of the ethylene precursor, aminocyclopropanecarboxylate (ACC), and a compound metabolized into ethylene, ethephon. For ACC, 5 mL of 1mM dilution (with distilled water) was sprayed every three days. For ethephon 50 mL of 0.1% dilution (with distilled water) was treated into planted soil every two days. For each treatment, each 5 control and treated samples were assessed, and they mostly consistent phenotypic changes.

**Supplementary Text S1 Searching for ETR1-like orthologs in the Male *Cannabis sativa* genome**

We reassembled the publicly available male *Cannabis sativa* (cultivar Finola) genome from raw reads and attempted to identify the ETR (ethylene receptor) 1-like gene within the sequence. PacBio Continuous Long Read (CLR) raw reads (accession No: SRR7274799) were downloaded from NCBI, and assembly was performed using Canu (parameters used: batOptions=-dg 3 -db 3 -dr 1 -ca 500 -cp 50). A tblastn search for the ETR1 gene (accession No: XP_030485641) was conducted on the assembled results, and the search yielded one hit with over 99% identity and two hits with approximately 70% identity, all showing more than 50% identity. Upon investigating the genomic region where the hit with over 99% identity was located, it was found to correspond to the annotated ETR1 gene on the X chromosome of the *Cannabis sativa* RefSeq (cultivar pink pepper). On the other hand, the two hits with approximately 70% identity were both located in regions annotated as ERS (ethylene response sensor) 1 on chromosome 7 in RefSeq. The two separate hits were likely caused by the assembly of genomes derived from allelic variants on homologous chromosomes. Based on these results, the ETR1-like gene was only found at one location on the X chromosome and nowhere else.

**Supplementary Text S2 A hypothetical ethylene receptor inhibitor, STS, properly acted for ethylene signal inhibition in *Humulus* species.**

Total RNA was extracted from early flower primordia of the STS-treated and control female *H. japonicus* (biological replicates *N*=5 for each), to prepare mRNA-seq libraries, as described (see Methods section). mRNA-seq reads were mapped to the genome of *H. japonicus* with BWA v0.7.17 to detect normalized expression level (RPKM). DESeq2 v1.44 (68) was used to screen the differentially expressed genes (DEGs) with the threshold of $P$ value < 0.05. As the result, we detected 294 and 971 genes up- or down-regulated in the STS-treated samples (table S8). The down-regulated genes included many genes reminiscent of ethylene signaling and ethylene response, which was supported by a GO enrichment analysis with TBtools v7.37.1 (69). The genes down-regulated in the STS treatment statistically enriched genes with GO terms of multiple “ethylene response” categories (table S9), and also defense-related genes, which is also often associated with ethylene signaling. Together, they indicated that a hypothetical ethylene receptor inhibitor, STS, could properly act for ethylene signal and response inhibition in *H. japonicas*.

**Figure S1**

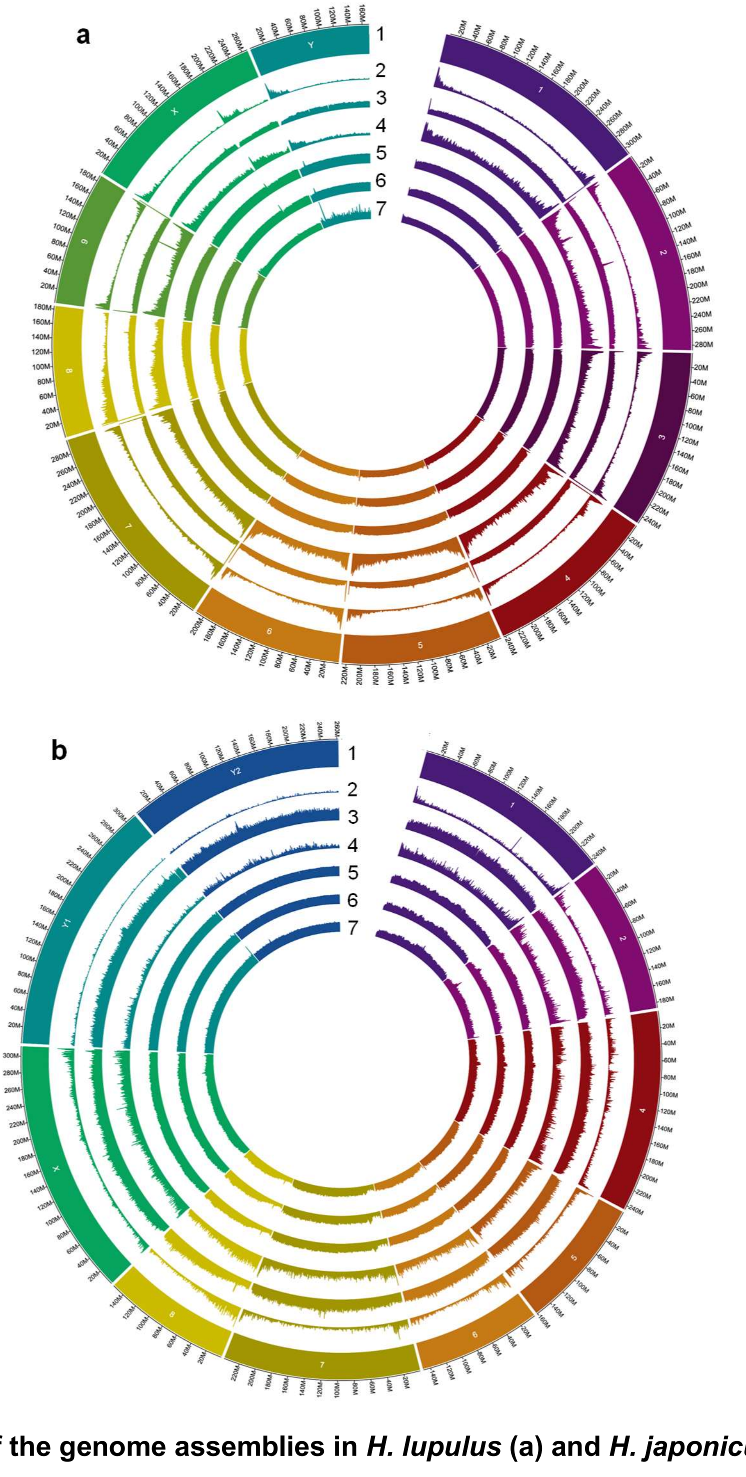


**Figure S1 Overview of the genome assemblies in *H. lupulus* (a) and *H. japonicus* (b)**

The same chromosome numbers were assigned to syntenic chromosomes in *H. lupulus* and *H. japonicus*. Due to the X-autosome fusion in *H. japonicus*, this species lacks Chr. 3, and its X is composed of parts syntenic with the *H. lupulus* X and its Chr. 3, as explained in Fig. 2. Layer 1: chromosome names, 2: Gene density, 3: LTR-type transposable element (TE) density, 4: non-LTR-type TE density, 5-7: DNA methylation levels in young male leaves (5: CGN, 6: CHG, and 7: CHH).

### Figure S2

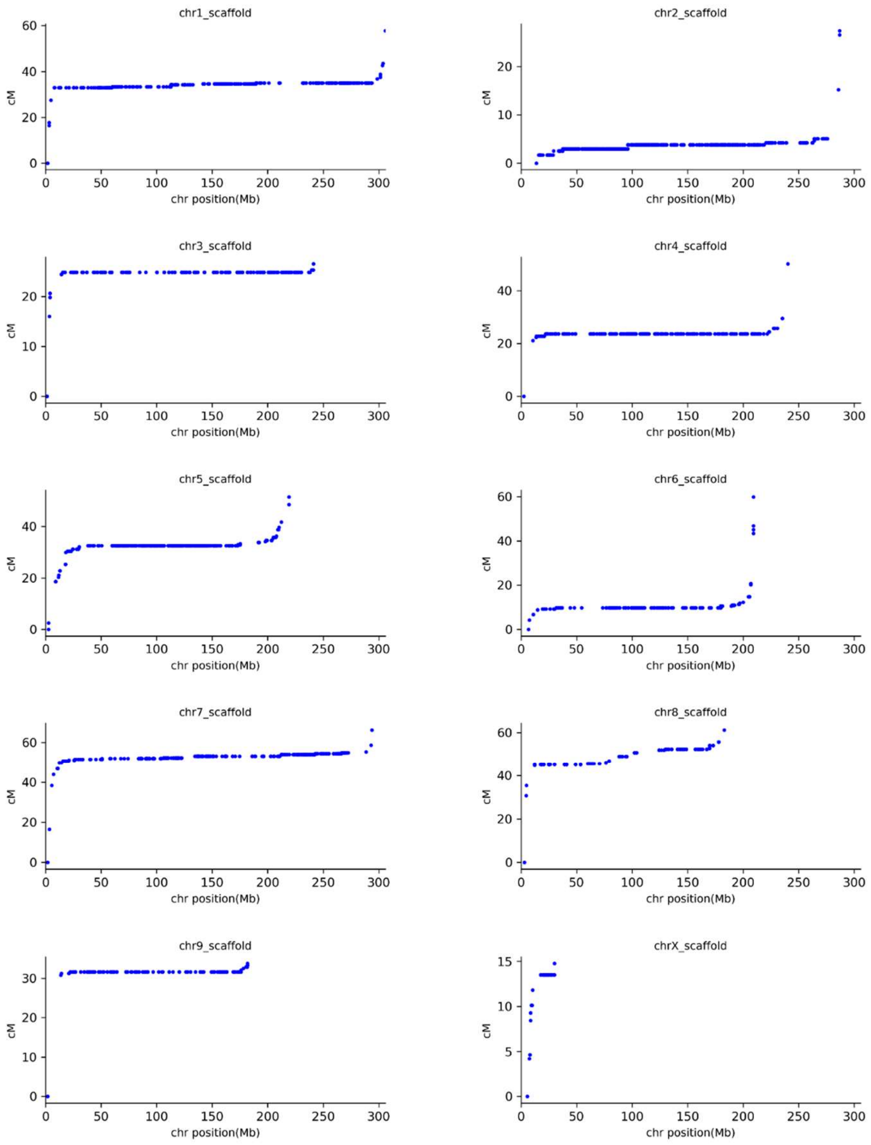


**Figure S2 Marey maps showing the relationship between the physical and genetic maps in a *H. lupulus* F1 population**

Male meiotic genetic map positions (y axes) of markers mapped in each chromosome, with marker physical positions in the x axes). Most chromosomes have extensive recombinationally inactive putatively pericentromeric regions. Recombination between the X and Y chromosomes was detected in the small recombinationally active regions, which occupies 0-35Mbp of the X assembly. Genes in this pseudo-autosomal region (or PAR) displayed allelic SNPs, indicating that the region is not degenerated, unlike the rest of the Y, in which many X-linked genes are hemizygous in males (see the main text). We also found some hemizygous Y-specific markers; consistent an absence of recombination, causing isolation between the Y and X, such Y-specific variants showed no recombination in MSY in our mapping population.

**Figure S3**

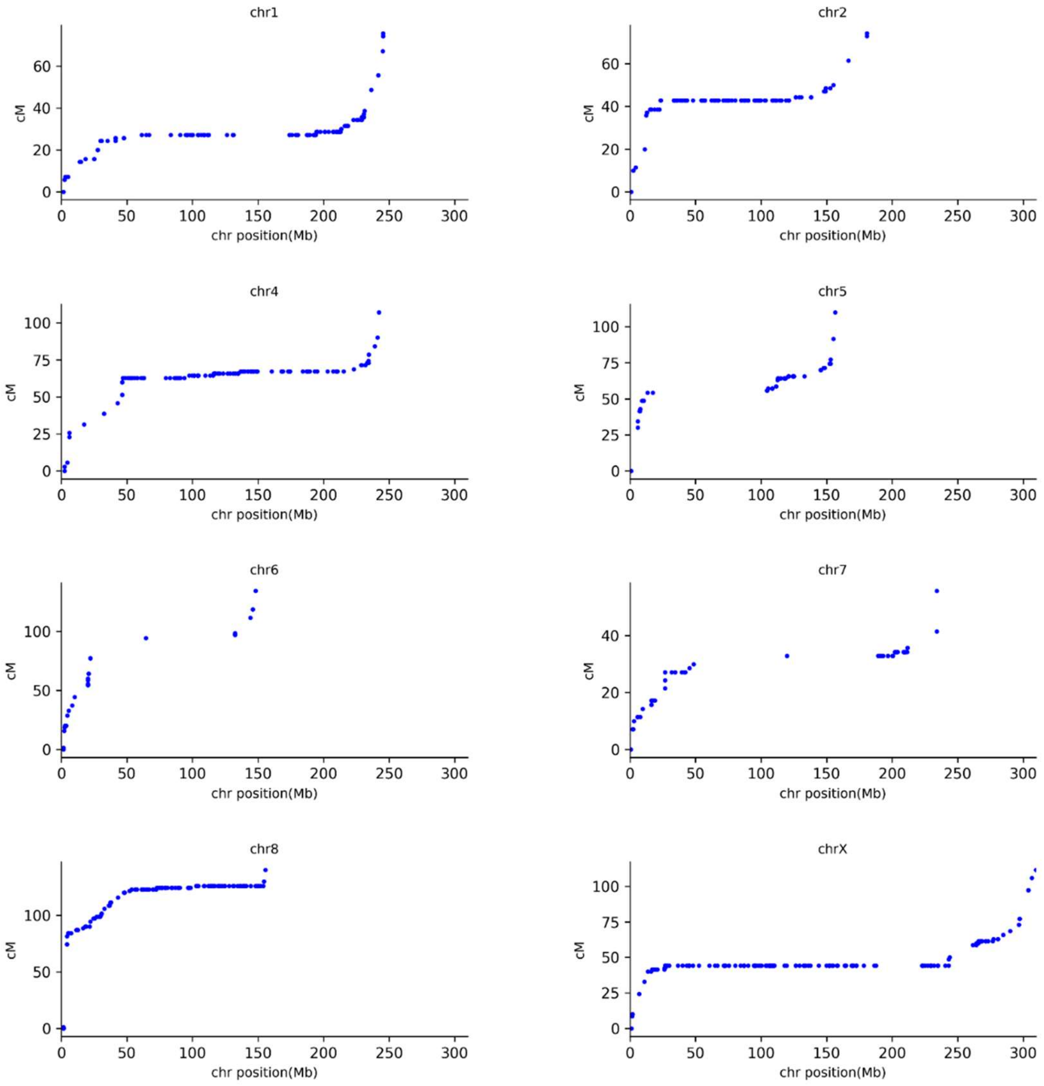


**Figure S3 Marey maps showing the relationship between the physical and genetic maps in a *H. japonicus* F1 population**

The axes are as in Figure S2, although genetic map is in female meiosis for the X chromosome, as male showed no recombination between X and Y chromosomes. Similar to the results from *H. lupulus*, most chromosomes have large recombinationally inactive regions. In this species, no allelic SNPs were shared between the X- and Y-linked sequences, suggesting that the Y is more degenerated than in *H. lupulus*. Recombination could be therefore be studied only in female meiosis. Although X-Y recombination could not be estimated, the evidence from the degeneration just described strongly suggests that it is absent throughout both parts of the large fused X-linked region. In other words, the former chromosome 3 is at least largely, and possibly entirely, non-recombining.

**Figure S4**

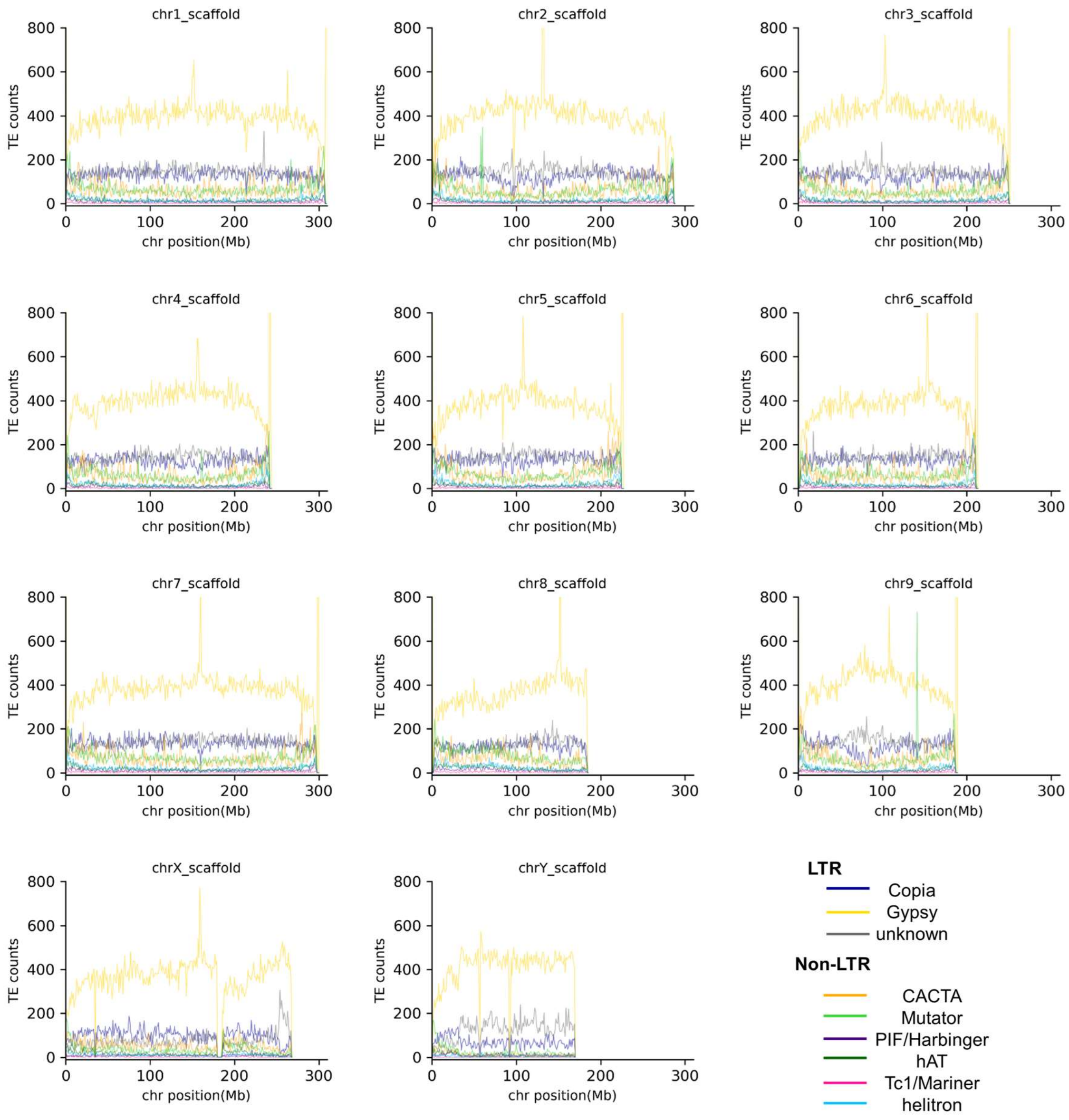


**Figure S4 Distribution of transposable elements in the assembled *H. lupulus* chromosomes**

The y axes show counts of three LTR-type classes (*Copia*, *Gypsy*, and *unknown*) and six classes of Non-LTR-type TEs (*CACTA*, *Mutator*, *PIF/Harbinger*, *hAT*, *Tc1/Mariner*, and *Helitron*) in 1Mb bins. Throughout the genome, the LTR class elements are most abundant. The chromosome ends tend to have smaller TE abundances.

**Figure S5**

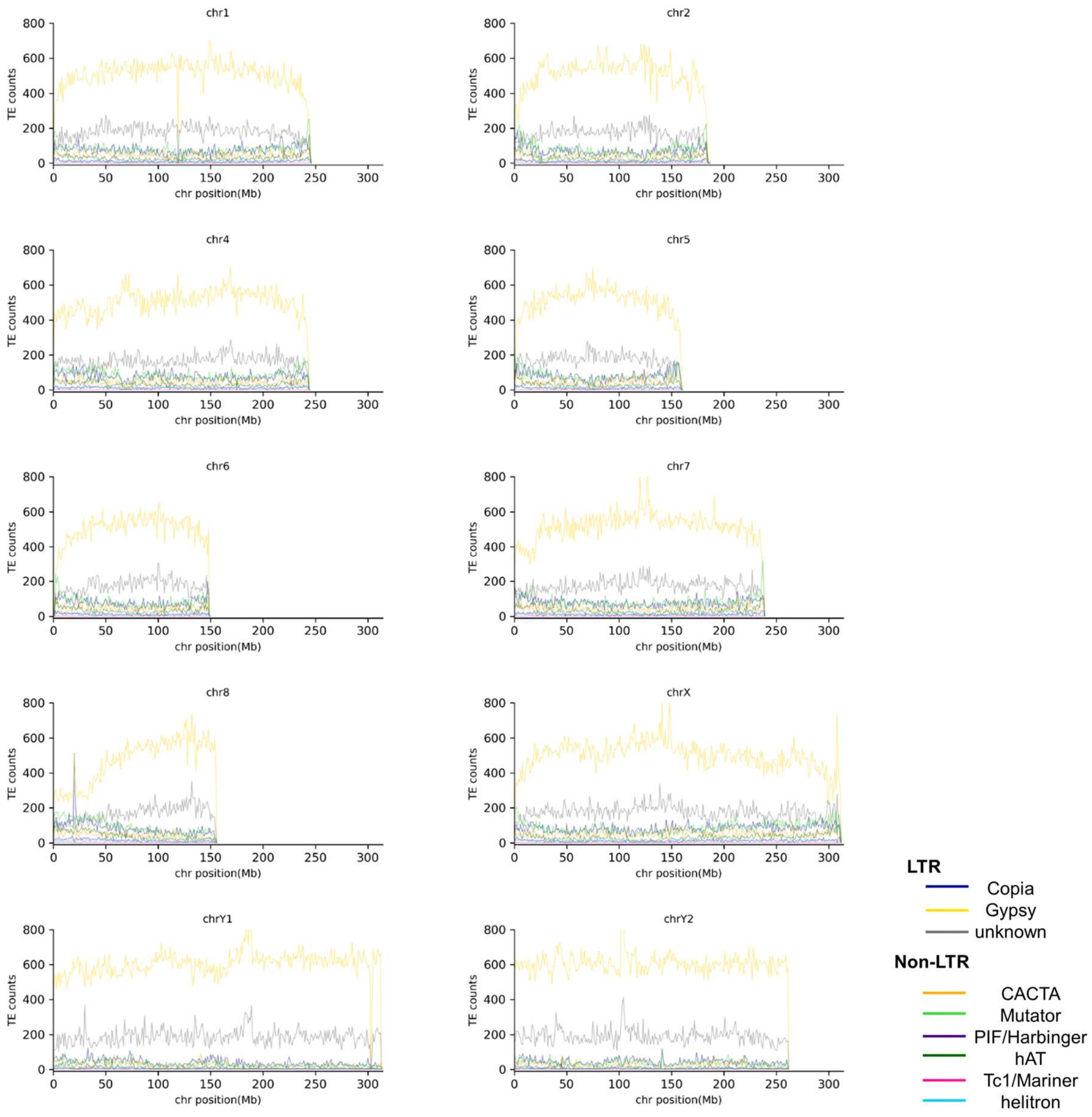


**Figure S5 Distribution of transposable elements in the assembled *H. japonicus* chromosomes**

The pattern is similar to that in *H. lupulus* (Figure S4).

**Figure 6**

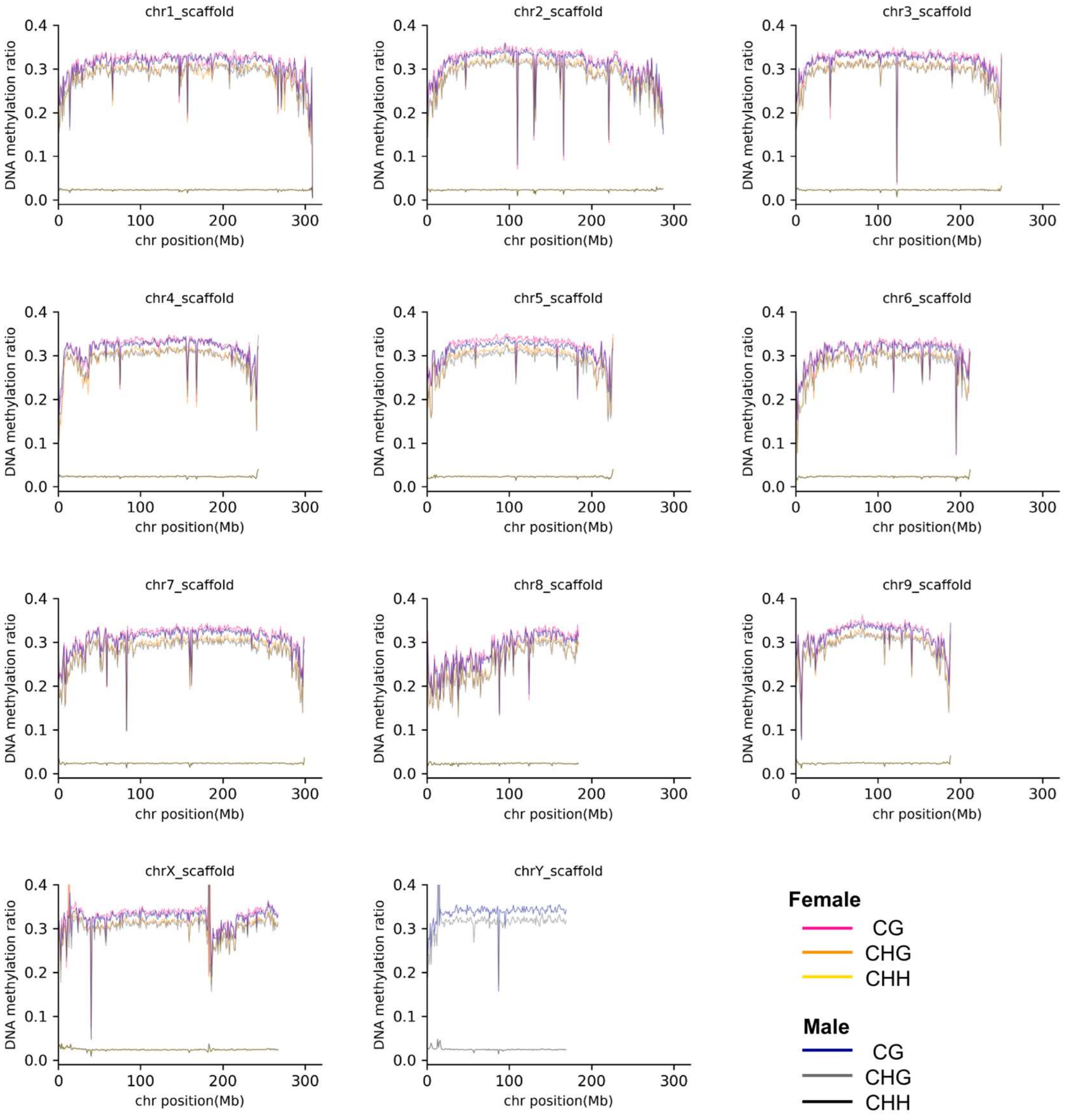


**Figure S6 Genome-wide DNA methylation levels in *H. lupulus***

DNAme levels in the three contexts, CGN, CHG, and CHH, in young leaves from male and female siblings in 1Mb bins. Chromosome ends tended to show lower DNA methylation levels, corresponding with the regions of high TE abundance (fig. S4).

**Figure S7**

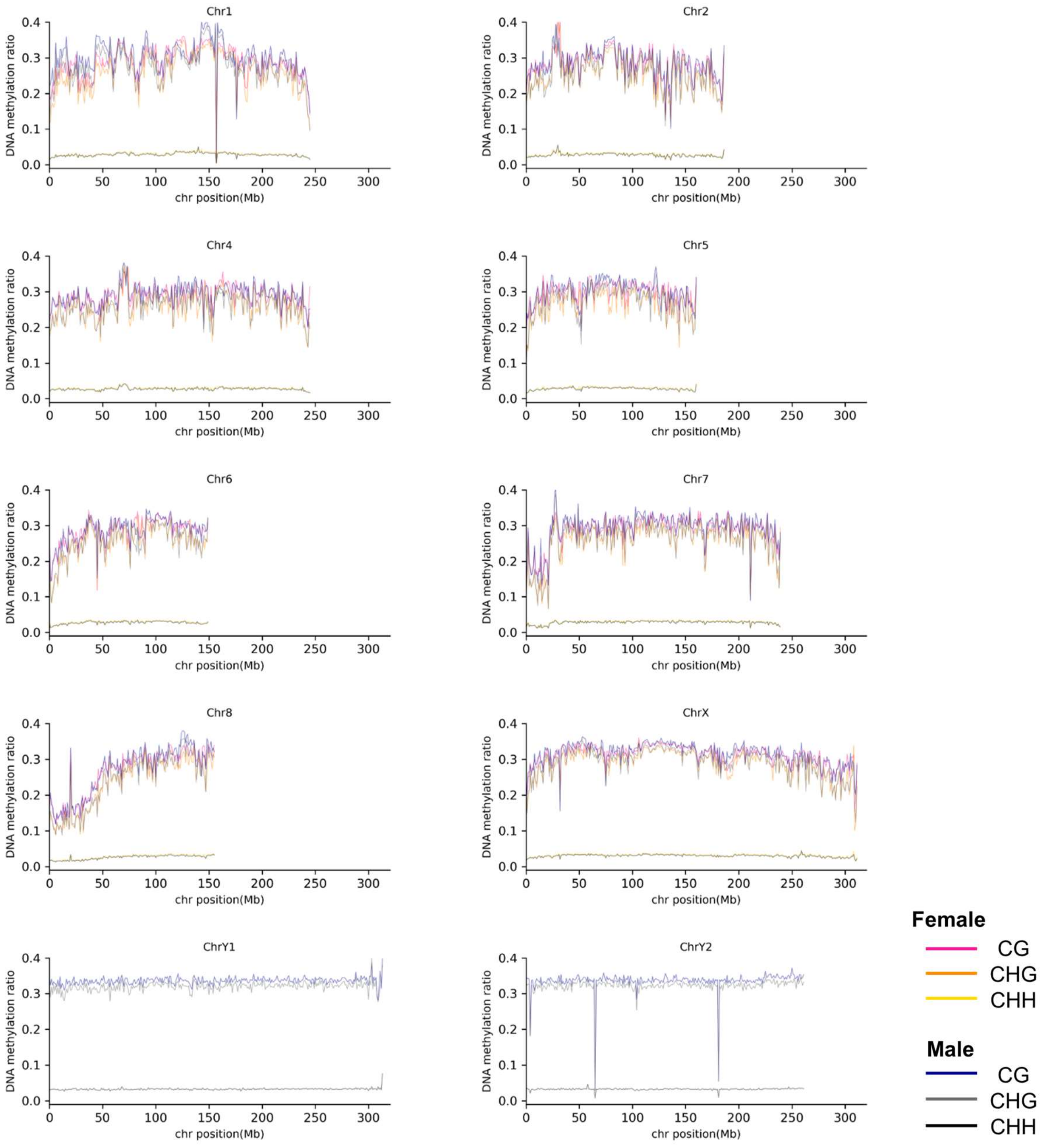


**Figure S7 Genome-wide DNA methylation levels in *H. japonicus***

DNAme levels in all three context (CGN, CHG, and CHH) in male and female young leaves (from a siblings) per 1Mb bin were plotted. Synchronized to the pattern of TE accumulation (fig. S5), chromosome both ends tended to show lower DNA methylation levels.

**Figure S8**

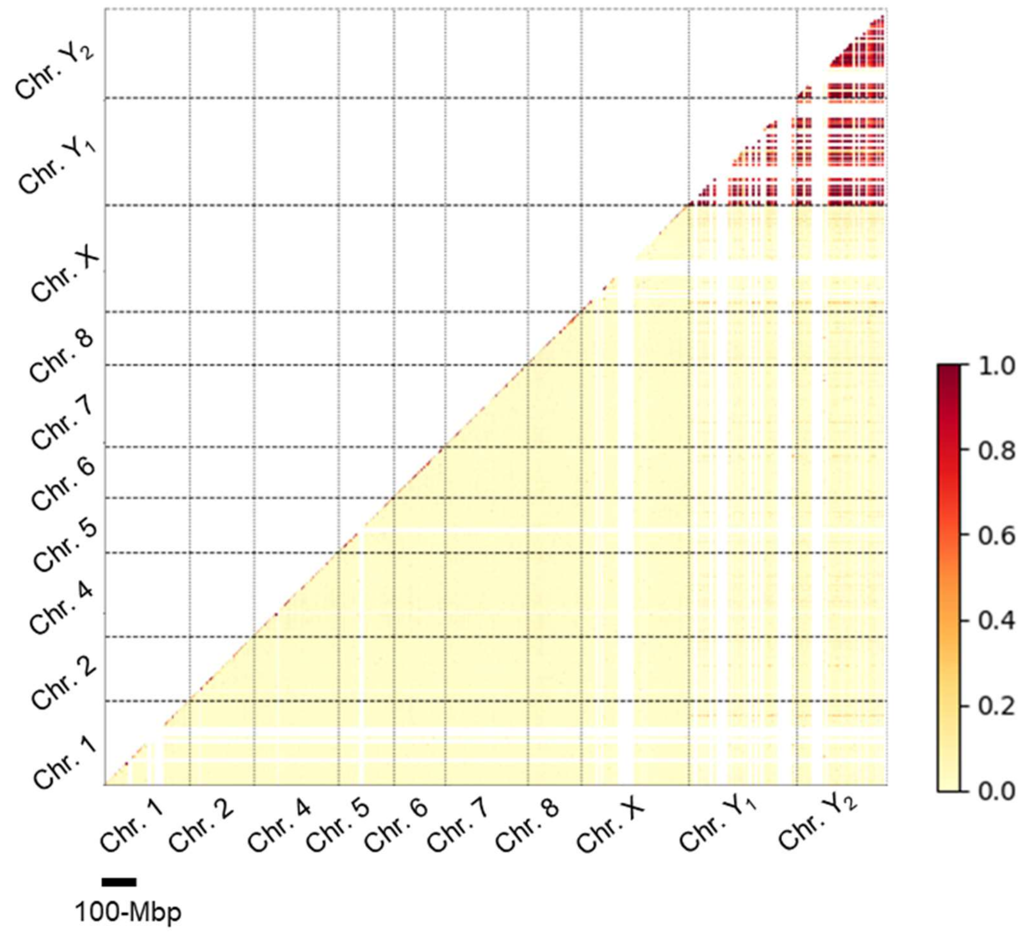


**Figure S8 Linkage disequilibrium (LD) heat map in a natural population of *H. japonicus***

Pairwise $r^2$ values between SNPs in different positions in the assembly are shown below the diagonal. In contrast to the autosomes and X chromosome, SNPs in sequences assembled on the $Y_1$ and $Y_2$ chromosome exhibited high values, both within these chromosomes and between them, consistent with the expected co-segregation of the two chromosomes.

**Figure S9**

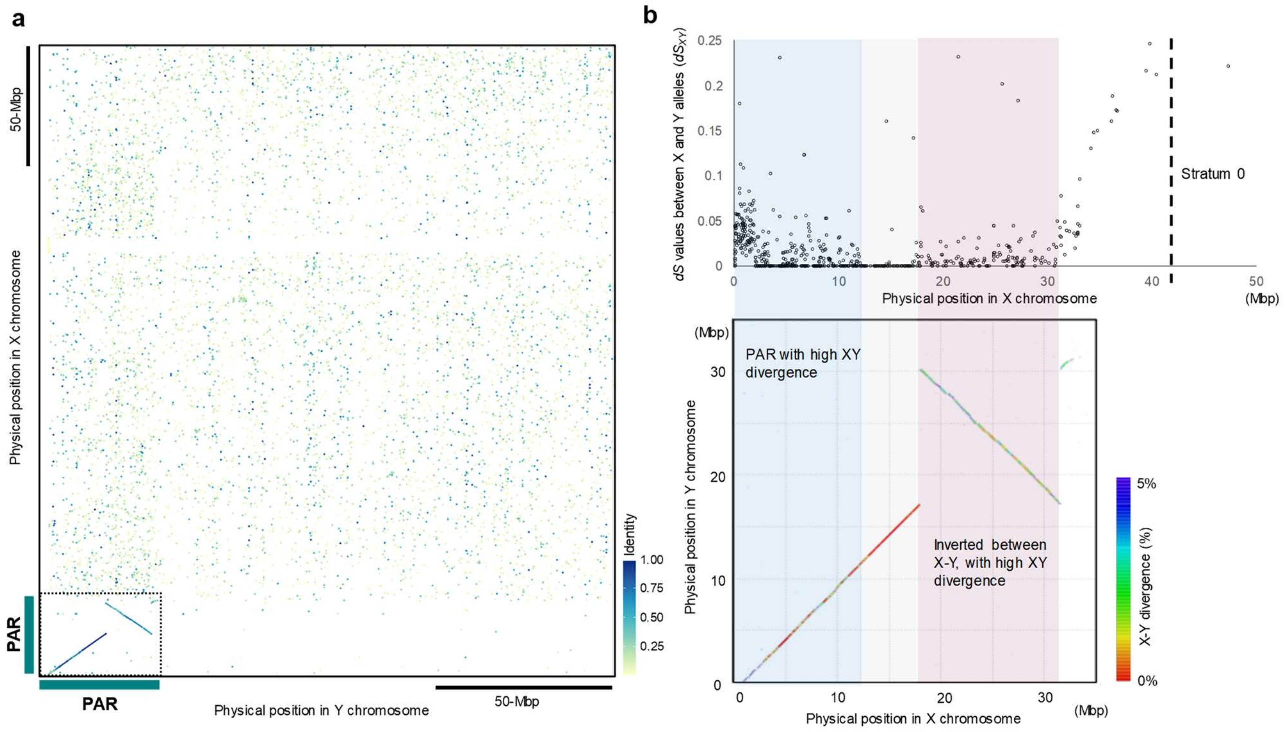


**Figure S9 Organisation of the *H. lupulus* X and Y chromosomes and the PAR**

**a**, Sequence-based synteny dot plot between this species' entire X and Y chromosomes. **b**, Only the approximately 30-35Mbp recombinationally active region (see fig. S2), the likely PAR, exhibits clear long syntenic regions, but after the initial ~18 Mb, a long inversion is present. Divergence estimates for synonymous sites ($dS_{XY}$ values) are shown across these two parts. The left-hand sub-region of the non-inverted region (indicated by the blue box) includes predominantly high values (often up to 5%), while the right-hand inverted (inverted, pink box) region has intermediate values that are consistently higher than those in the middle of the entire region, whose $dS_{XY}$ values are consistently very low, suggesting that only this smaller set of genes are PAR genes.

**Figure S10**

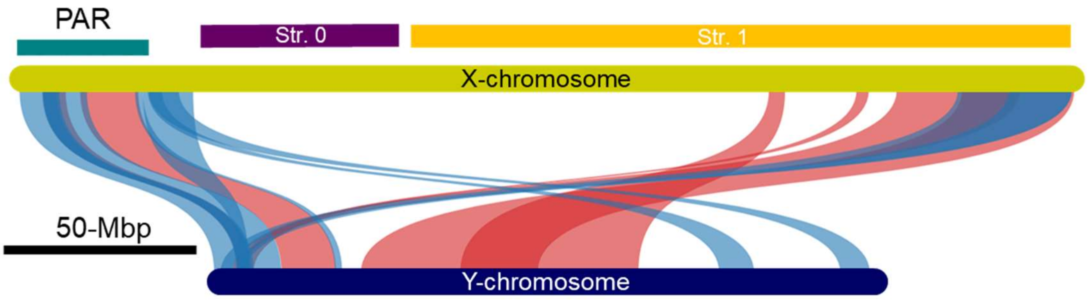


**Figure S10 Synteny analysis between the *H. lupulus* X and Y chromosomes in based on the order of genes.**

The analysis used MCScanX with a low threshold value (e-value = $1e^{-10}$). Based on genes identified in the X chromosome assembly, the analysis suggests that about half of Stratum 1 is still syntenic with the Y. Blue and red bands respectively indicate the same and opposite directions in the two chromosome assemblies.

**Figure S11**

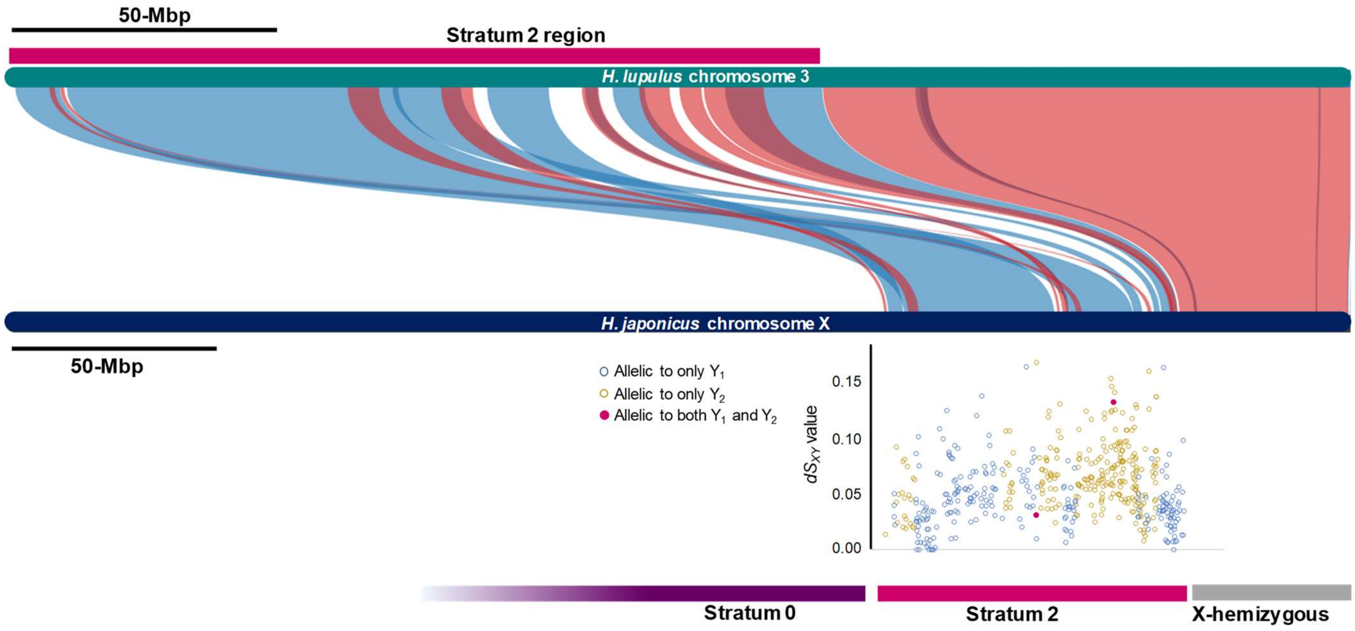


**Figure S11 Frequent rearrangement in Stratum 2 in *H. japonicus***

The X-hemizygous region in the right arm of the enlarged *H. japonicus* X exhibits clear synteny to the *H. lupulus* chromosome 3 (fig. S13). However, in the Stratum 2 region, synteny is observed only in discontinuous blocks that partially correspond to $Y_1$ or $Y_2$, with different $dS_{XY}$ values. Note that the $dS_{XY}$ values for individual genes have high variance, and the values fluctuate too highly for a change-point test to succeed. Blue and red bands respectively indicate the same and opposite directions in the two chromosome assemblies.

**Figure S12**

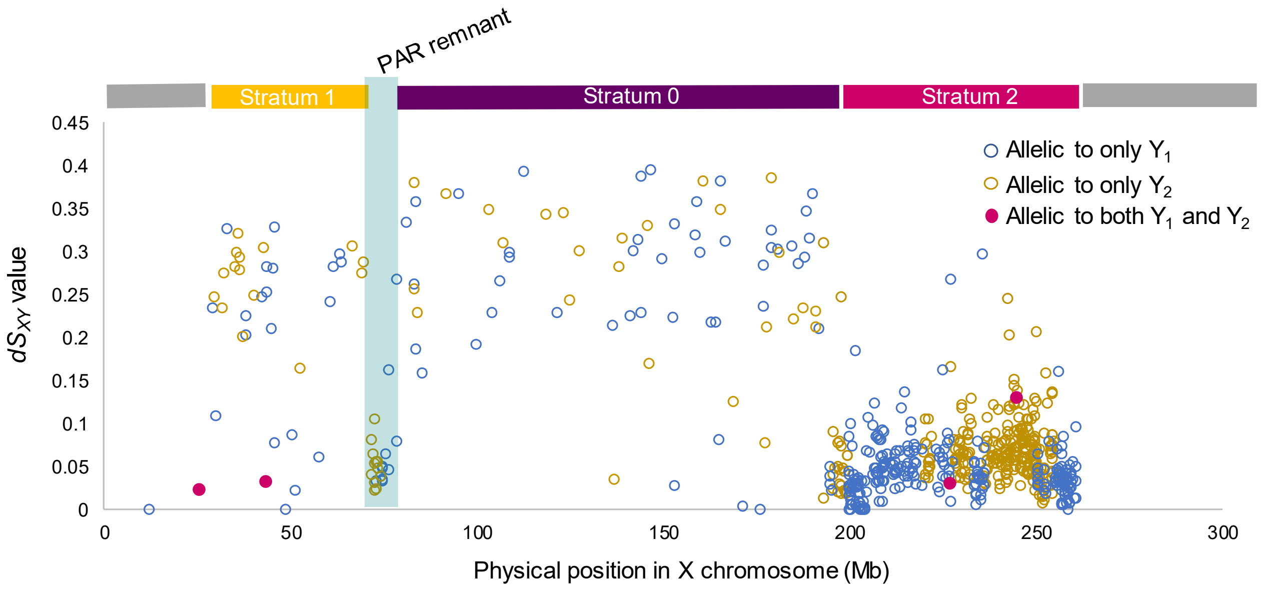


**Figure S12 Relationships between X and $Y_1$/$Y_2$ chromosome sequences in *H. japonicus***

The y axis shows $dS_{XY}$ values of inferred gametologous gene pairs (with genes annotated on both the X and an apparently Y-linked allele), with their X chromosome positions on the x axis. As expected, the Y-linked sequences of most such X-linked genes are on $Y_2$ (derived from Hl3) with only a few on $Y_1$ (derived longer ago, from the ancestral X, whose Y is profoundly degenerated). However, the Hl3 genes are not confined to the $Y_2$ as would be expected, and HlX ones are not confined to the $Y_1$; instead, gametologs from each set are found on both $Y_1$ and $Y_2$, with HlX-derived genes in Strata 0 and 1 apparently randomly distributed between the two Ys, and Hl3-derived genes in Stratum 2 having a mosaic distribution, with (very unexpectedly) many found only in $Y_1$. A few genes (indicated by the 4 magenta dots) are shared by both $Y_1$ and $Y_2$. The mosaic pattern observed in $Y_2$ suggests rearrangements in which segments of $Y_1$ moved to $Y_2$ (or were perhaps duplicated onto chromosome 3 before it became $Y_2$), while a few $Y_2$ genes moved to $Y_1$.

**Figure S13**

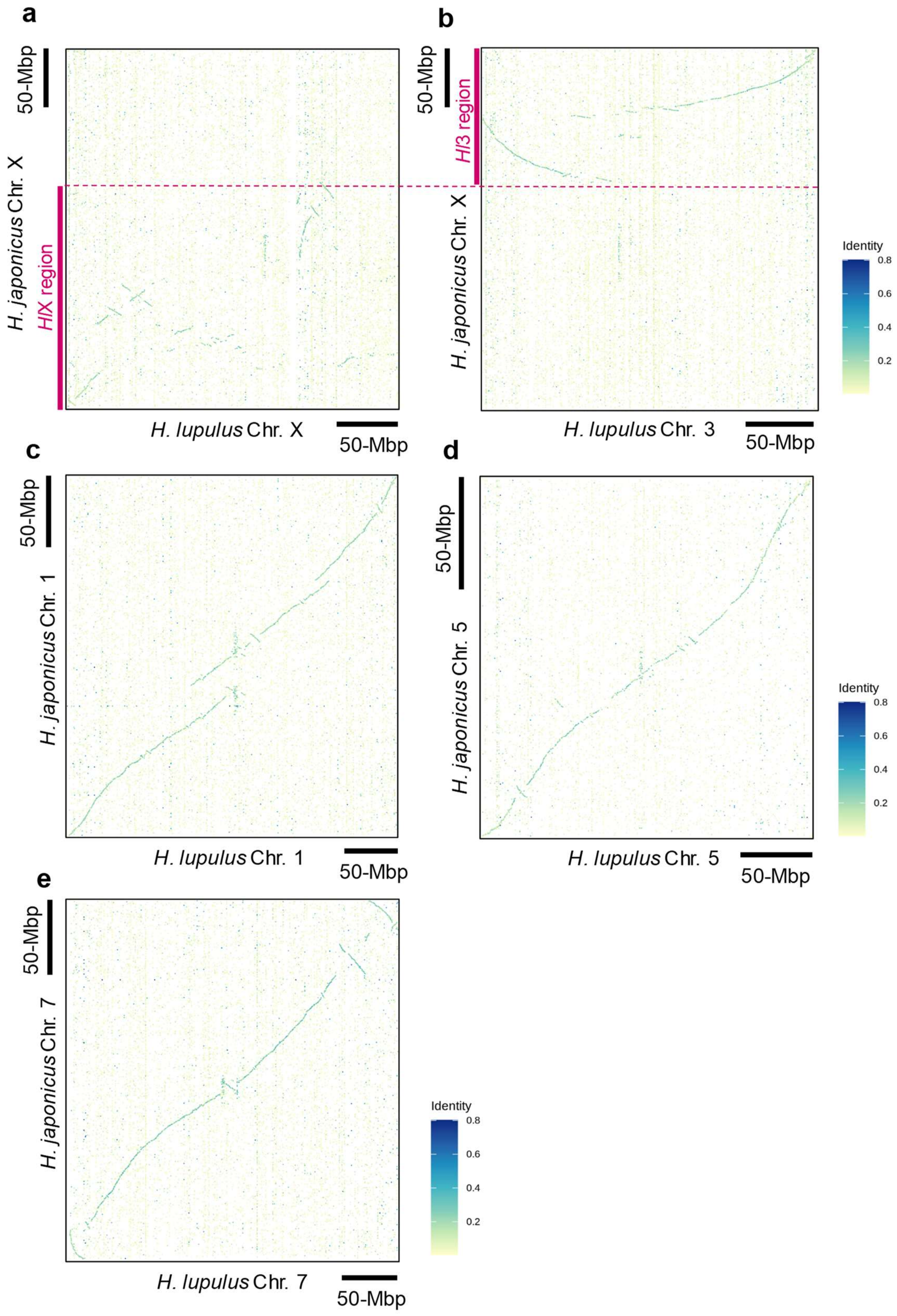


**Figure S13 Sex chromosome-specific rearrangements**

Synteny plots between *H. lupulus* and *H. japonicus*, in three representative autosomes and their X chromosomes. Highly disrupted synteny in the X chromosome (**a, b**), than in the autosomes (**c**: Chr. 1, **d**: Chr. 5, **e**: Chr. 7). The X chromosome in *H. japonicus* is consitituted of *Hl*X and *Hl*3, and *Hl*3 region showed less disrupted synetny (against Chr. 3, an autosome in *H. lupulus*) than *Hl*X. This is presumably due to a comparison between X (in *H. japonicus*) and autosome (in *H. lupulus*), and also due to recent conversion from an autosome to a part of Chr. X in *Hl*3.

**Figure S14**

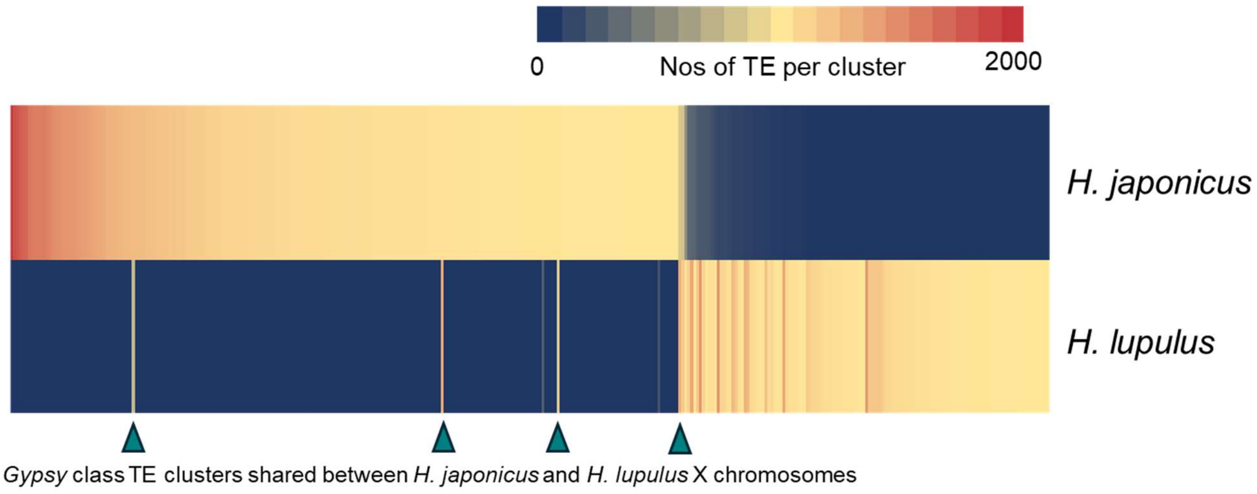


**Figure S14 *Gypsy*-class TE clustering in X chromosome of *H. lupulus* and *H. japonicus***

Clustering of *Gypsy*-class TE identified in the X chromosomes of *H. lupulus* and *H. japonicus*, with allowance of net divergence = 0.2 (which is fundamentally higher than the *H. lupulus-H. japonicus* interspecific silent divergence: 0.128). Hence, most of the TE were thought to burst after the divergence of *H. lupulus* and *H. japonicus*, with only a few exceptional clusters indicated by arrow heads.

**Figure S15**

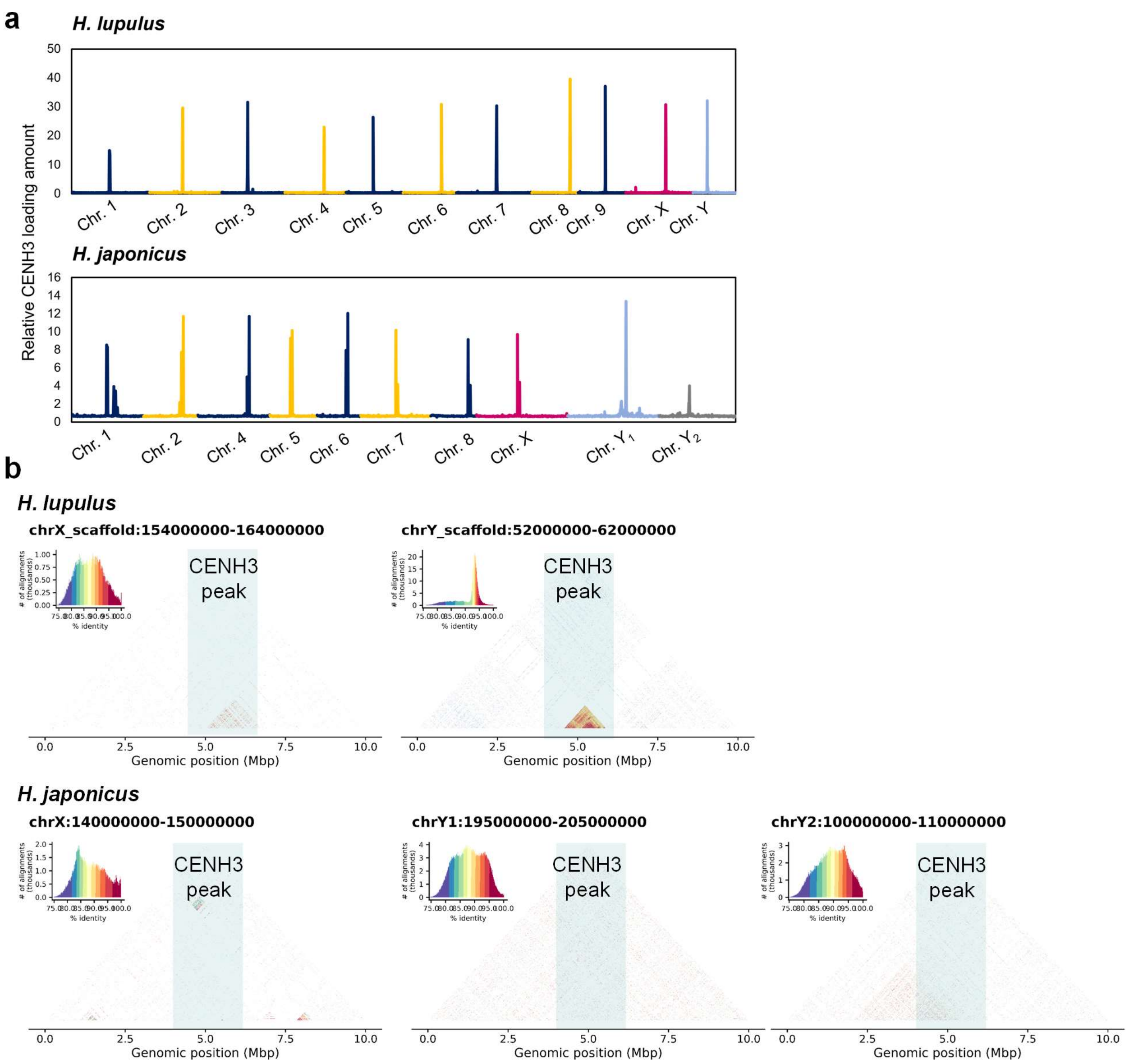


**Figure S15 Definition of centromeric regions and centromeric repeats in sex chromosomes**

**a**, Detection of peaks of ChIP-seq analysis with CENH3 antibody in *H. lupulus* and *H. japonicus.* In all chromosomes, CENH3 loading amount exhibited clear single (or partially double) peaks, which can be defined as centromeric regions. **b**, Tandem repeat visualization in the CENH3-peaked region in sex chromosome of *H. lupulus* and *H. japonicus*. *H. lupulus* showed mature tandem repeats in the centromeric regions in both X and Y chromosomes. On the other hand, *H. japonicus* showed no clear tandem repeats specific to the centromeric regions.

**Figure S16**

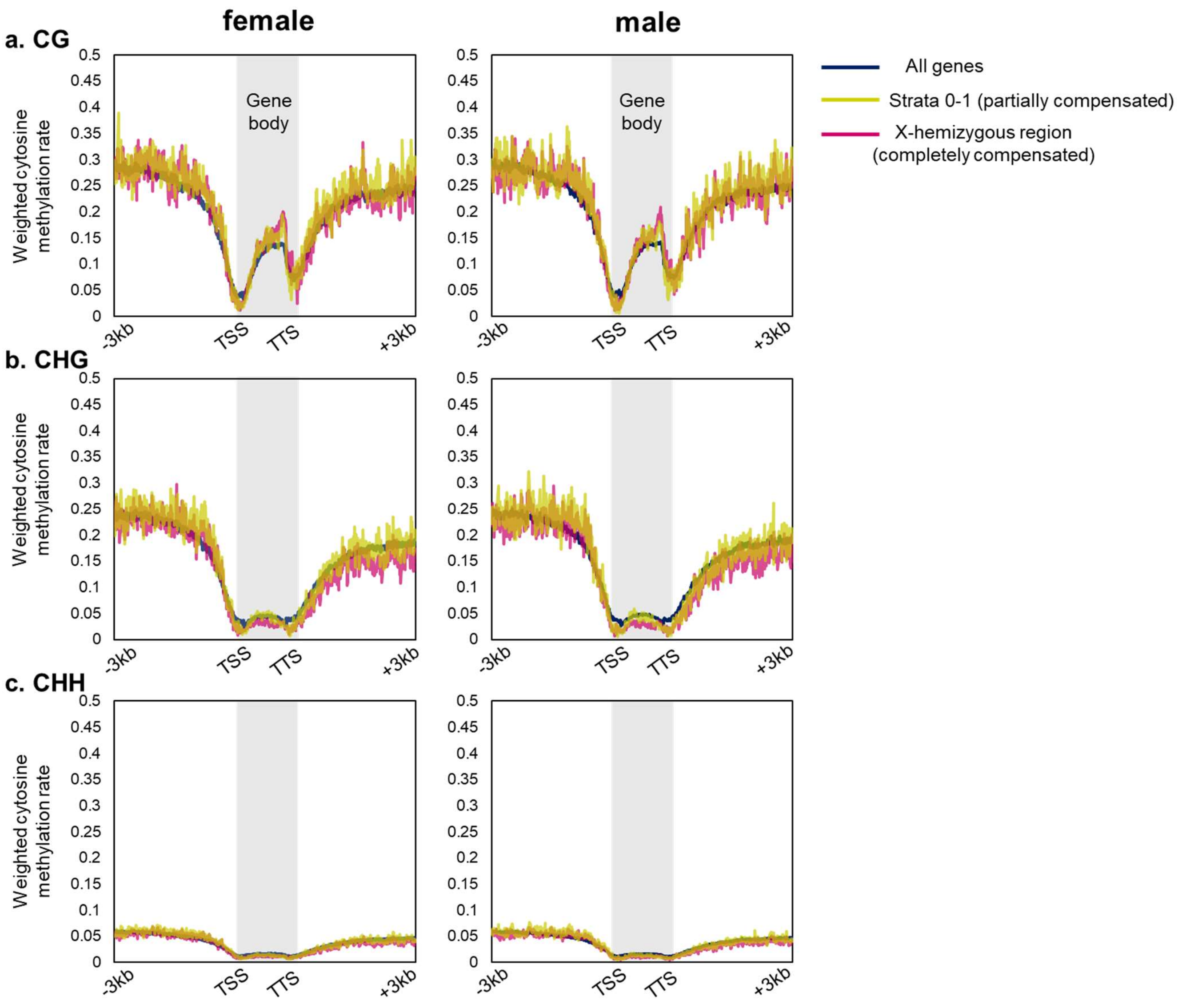


**Figure S16 DNA methylation status is not involved in dosage compensation of *H. japonicus***

Transition of DNA methylation levels surrounding the genes in the whole genome, partially compensated regions (strata 0-1), and completely compensated regions (X-hemizygous regions). In all three context (CG, CHG, CHH), no substantial differences were detected among the regions and also between male and female.

**Figure S17**

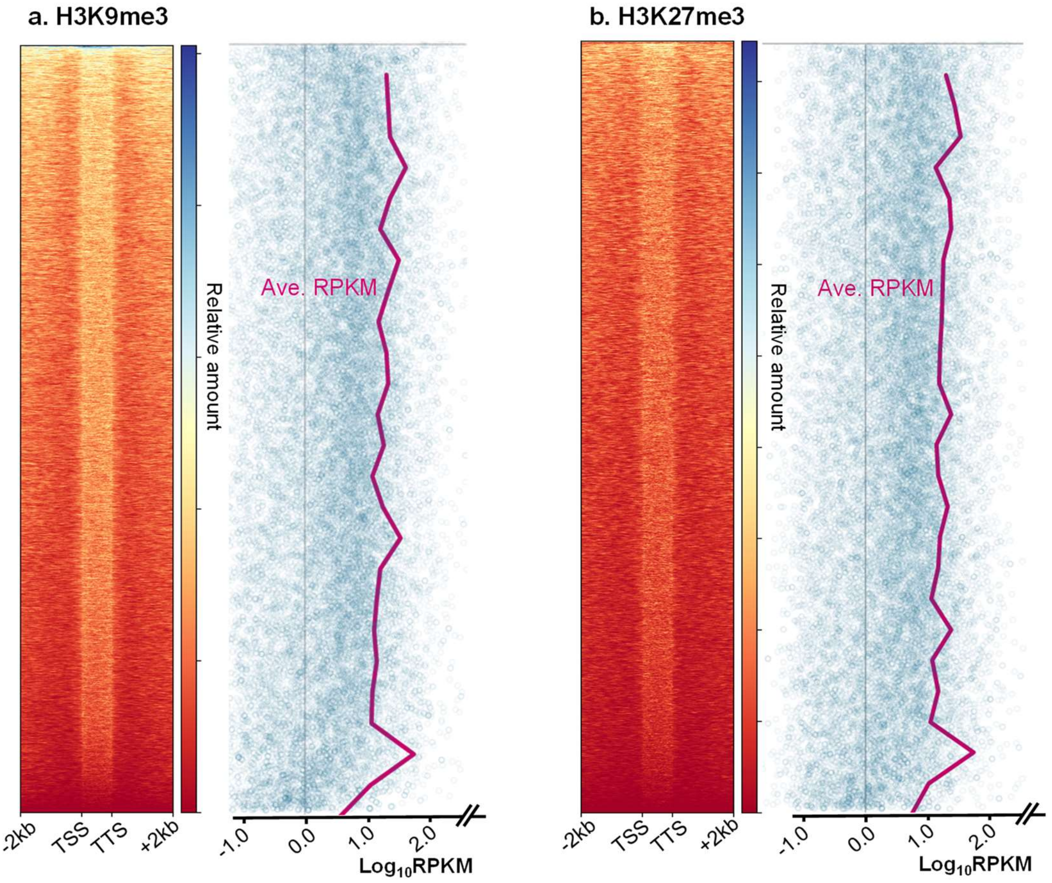


**Figure S17 Clustering of H3K9me3 and H3K27me3 modification patterns in the whole genes, and their correlations with expression levels.**

Modification patterns of H3K9me3 (**a**) and H3K27me3 (**b**) around genes in the *H. japonicus* genome. The genes' expression levels (averaged values of 4 female leaves) are neither positively nor negatively correlated with the pattern, in either histone methylation context (in female).

**Figure S18**

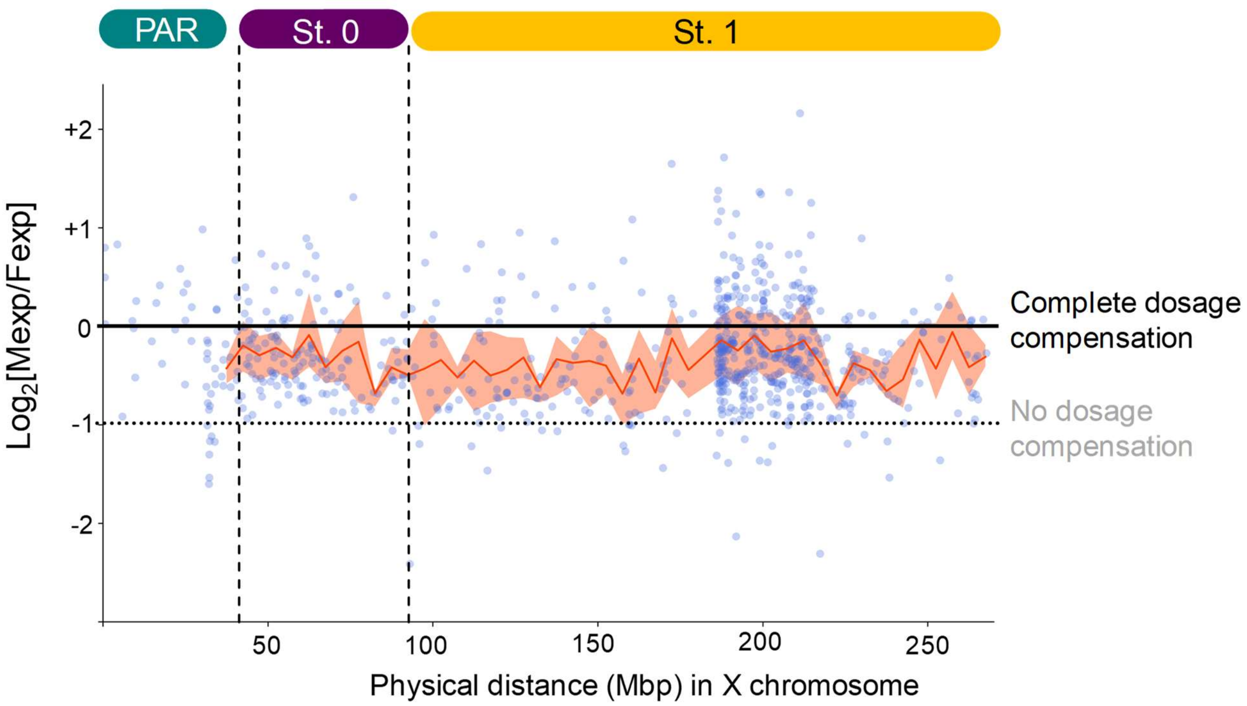


**Figure S18 Dosage compensation in *H. lupulus* X chromosome**

Plots of male-female expression bias ($log_2$[male-expression/female-expression]) in the X-specific genes of *H. lupulus*. Orange lines indicate the averaged values in 5-Mbp bin (with SD values indicated by ribbons). Throughout the Stratum 0-1, partial compensation was observed, regardless of the oldness of the strata and genomic positions/contexts. This tendency is contrast to the case of *H. japonicus* (Fig. 3).

**Figure S19**

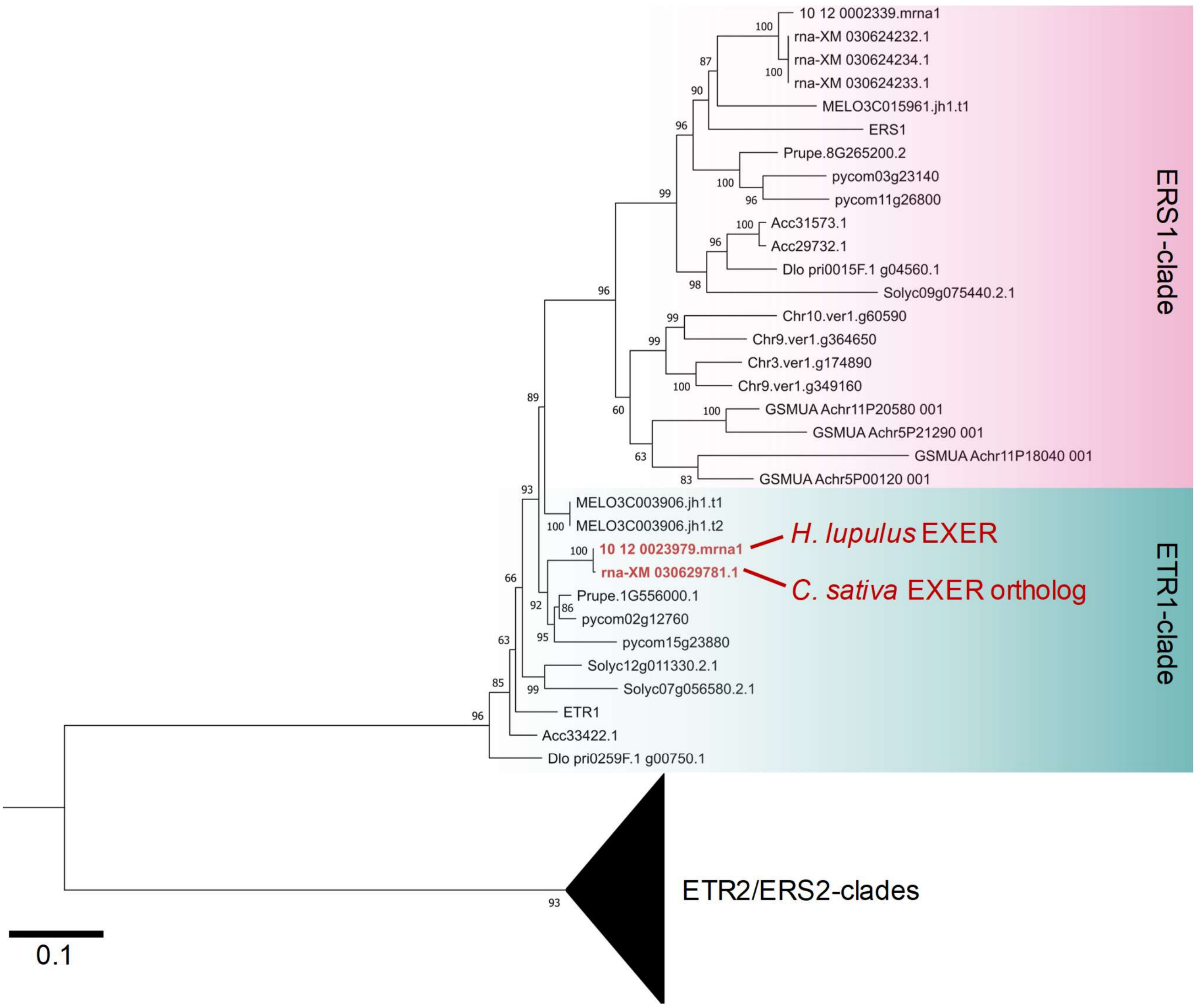


**Figure S19 Evolutionary topology of the ETR1/ERS1-clade ethylene receptors**

Evolutionary topology of the ETR1/ERS1-clade ethylene receptors were constructed with ML-method, with ETR2/ERS2-clades as the outgroup. Bootstrap values (with 100 replicates) were given on each branch. ETR1 and ERS1 clades clearly separated with statistic support (bootstrap=96/100). ETR1 clade is basically monophyletic, except lineage-specific duplications, and includes the sex determinant candidate, EXER gene, in the genera *Humulus* and *Cannabis*. For prefixes, Chr: avocado (*Persea americana*), GSMUA: banana (*Musa accuminata*), MELO: melon (*Cucumis melo*), Prupe: peach (*Prunus persica*), pycom: European pear (*Pyrus commonis*), Solyc: tomato (*Solanum lycopersicum*), Acc: kiwifruit (*Actinidia chinensis*), Dlo: *Diospyros lotus*, rna-XM: *Cannabis sativa*, 10_12: *Humulus lupulus*.

**Figure S20**

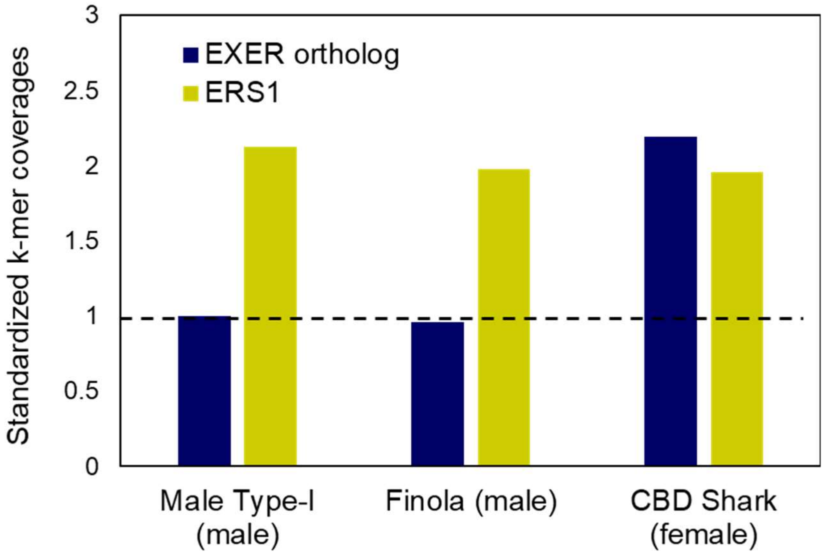


**Figure S20 Allele dosage of *EXER* in male and female *Cannabis sativa***

Standardized k-mer coverages in *EXER* and *ERS1* were calculated from the random gDNA sequencing data in two male and one female *C. sativa* (SRR10578258, SRR24187786, and SRR14857079, respectively). The dosage of *EXER* gene in Male Type-I was defined as “1”. This result suggested that *EXER* is X chromosome-specifically conserved, not only in *Humulus* but also in *Cannabis*.

### Figure S21

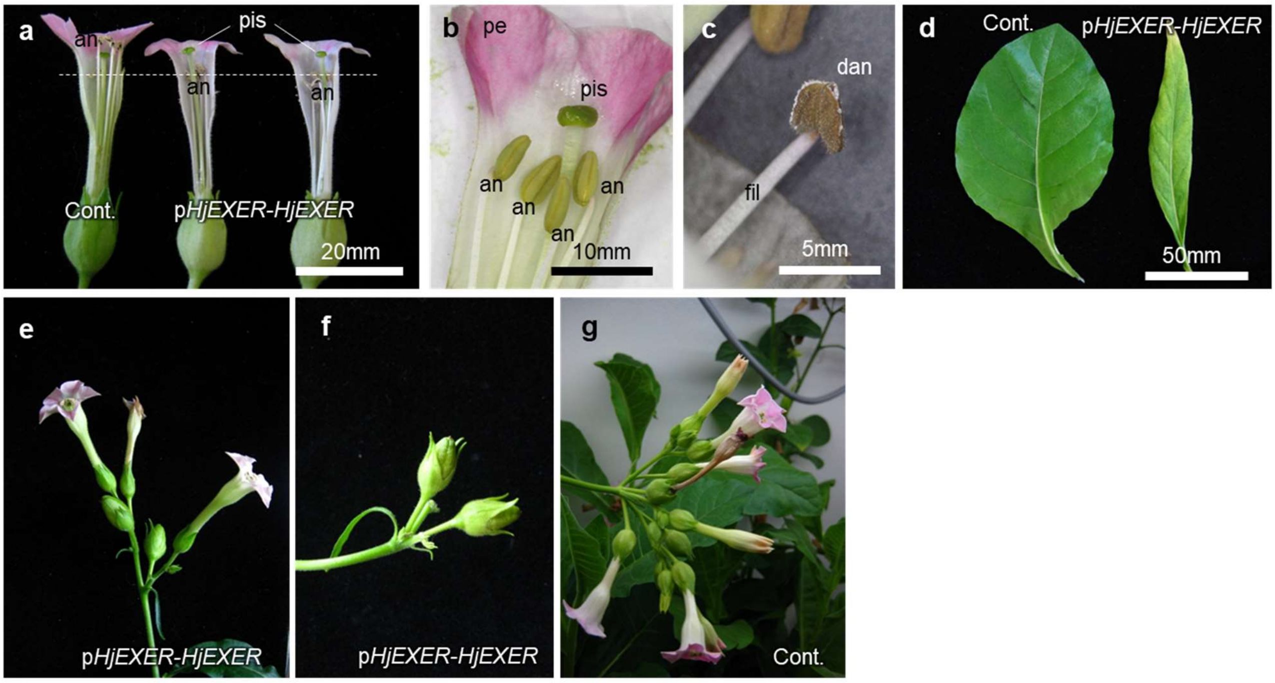


**Figure S21 Various phenotypes in *N. tabacum* with ectopic expression of the *HjEXER* gene**

**a**, *N. tabacum* plants transformed with *HjEXER* under the control of native promoter (*pHjEXER-HjEXER*) frequently exhibited fundamentally shorter androeciums (or feminization) than control plants. an: anther, pis: pistil. **b-c**, In *pHjEXER-HjEXER* plants, anthers are often defected or unmatured. dan: defected anthers, fil: filament, pe: petals. **d**, *pHjEXER-HjEXER* plants often exhibited narrow leaves. **e-g**, *pHjEXER-HjEXER* plants (**e-f**) exhibited substantially less flowers per inflorescence than control plants (**g**). These phenotypes (narrow leaves and less flowers per inflorescence) are consistent with the feminized *N. tabacum* transformed with a HD-ZIP1 homeodomain-like feminization factor *MeGI* in *Diospyros* species (70).

**Figure S22**

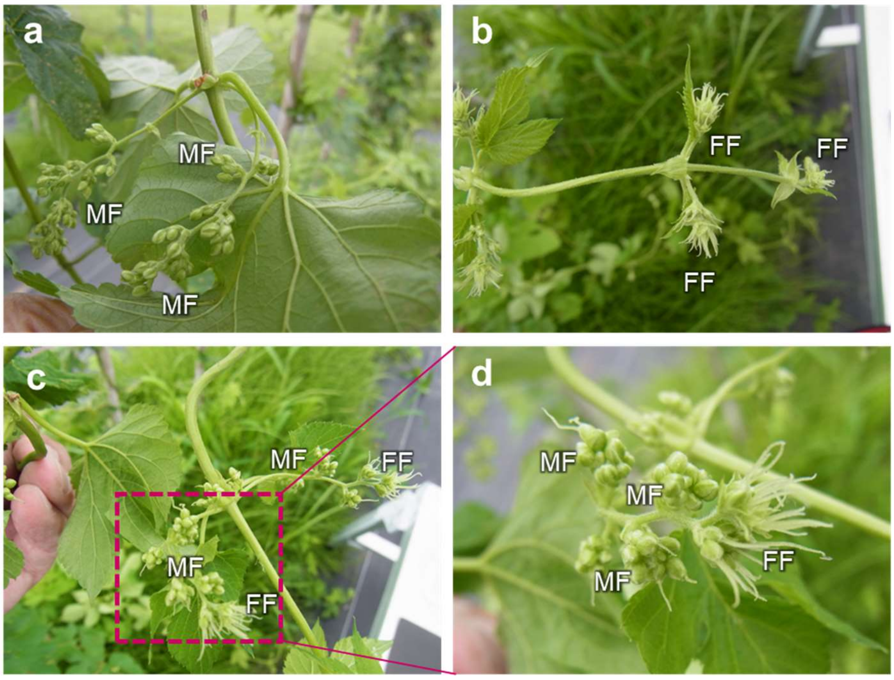


**Figure S22 Sex conversion with STS treatment in *H. lupulus***

**a**, Control non-treated male. **b**, Control non-treated female. MF: male flowers, FF: female flowers. **c-d**, Treatment with 2mM STS in genetically female *H. lupulus* plants resulted in formation of male flowers (or conversion into monoecy), consistent with the sex conversion in *H. japonicus* (Fig. 4). These results suggested that, in common in the genus *Humulus*, inhibition of the ethylene receptor function induces sex conversion from female to male.

### Table S1 Assembly statistics of *Humulus* genomes

| | | *Humulus lupulus* (Hop) | | | | *Humulus japonicus* |
|---|---|---|---|---|---|---|
| | | Isolate 10-12 | | Cultivar Saaz | | - |
| | | Male | | Female | | Male |
| Number of chromosomes | | 2n =20 (18 + X, Y) | | 2n =20 (18 + X, X) | | 2n = 17 (14 + X, $Y_1$, $Y_2$) |
| Assembly type | | Pseudo-haploid (haploid) | Haplotype-resolved (diploid) | Pseudo-haploid (haploid) | Haplotype-resolved (diploid) | Pseudo-haploid (haploid) |
| Genome assembly | Total length (bp) | 2,674,927,358 | 4,909,271,258 | 2,521,714,900 | 5,237,456,007 | 2,307,737,406 |
| | Number of sequences | 164 | 318 | 338 | 3,074 | 379 |
| | Largest contig length (bp) | 309,743,552 | 307,987,767 | 309,905,584 | 309,434,124 | 313,071,349 |
| | Number of N50 sequences | 5 | 9 | 5 | 10 | 5 |
| | Length of N50 sequences (bp) | 250,774,493 | 250,495,383 | 247,167,054 | 248,861,128 | 245,096,462 |
| | Total number of Ns (bp) | 1,000 | 6,000 | 4,500 | 129,000 | 22,200 |
| Genome evaluation | Complete | 98.15% | 98.24% | 98.02% | 98.28% | 98.20% |
| (BUSCO: eudicots_odb10) | Complete and single-copy | 94.80% | 0.06% | 91.10% | 4.56% | 92.30% |
| | Complete and duplicated | 3.35% | 97.64% | 6.92% | 93.72% | 5.80% |
| | Fragmented | 0.34% | 0.39% | 0.43% | 0.39% | 0.60% |
| | Missing | 1.50% | 1.38% | 1.55% | 1.33% | 1.20% |
| Gene prediction | | 26,490 | 49,594 | 25,979 | 52,194 | 26,320 |
| Gene evaluation | Complete | 98.54% | 98.84% | 98.75% | 98.88% | 97.54% |
| (BUSCO: eudicots_odb10) | Complete and single-copy | 91.75% | 4.77% | 95.36% | 0.99% | 89.17% |
| | Complete and duplicated | 6.79% | 94.07% | 3.40% | 97.89% | 8.38% |
| | Fragmented | 0.26% | 0.04% | 0.0% | 0.0% | 0.26% |
| | Missing | 1.20% | 1.12% | 1.25% | 1.12% | 2.19% |

**Table S2. Composition of telomeres and assembly gap in the *H. lupulus* cv. Saaz (female) haploid and diploid genomes.**

| **Haploid** | Telomere | | |
|---|---|---|---|
| Chromosome | 5' end | 3' end | Gaps (count) |
| chr1 | - | Detect | 0 |
| chr2 | - | Detect | 5 |
| chr3 | - | - | 2 |
| chr4 | Detect | - | 0 |
| chr5 | Detect | Detect | 1 |
| chr6 | - | Detect | 1 |
| chr7 | Detect | Detect | 0 |
| chr8 | Detect | - | 0 |
| chr9 | - | - | 0 |
| chrX | Detect | Detect | 0 |

| **Diploid** | Telomere | | |
|---|---|---|---|
| Chromosome | 5' end | 3' end | Gaps (count) |
| chr1_hap1 | - | Detect | 4 |
| chr1_hap2 | Detect | - | 0 |
| chr2_hap1 | - | - | 21 |
| chr2_hap2 | Detect | - | 36 |
| chr3_hap1 | - | - | 7 |
| chr3_hap2 | - | - | 5 |
| chr4_hap1 | - | Detect | 7 |
| chr4_hap2 | - | - | 4 |
| chr5_hap1 | - | Detect | 19 |
| chr5_hap2 | - | - | 23 |
| chr6_hap1 | Detect | - | 2 |
| chr6_hap2 | - | - | 1 |
| chr7_hap1 | Detect | - | 6 |
| chr7_hap2 | Detect | Detect | 5 |
| chr8_hap1 | - | - | 1 |
| chr8_hap2 | - | - | 0 |
| chr9_hap1 | - | - | 9 |
| chr9_hap2 | - | - | 9 |
| chrX_hap1 | - | Detect | 52 |
| chrX_hap2 | Detect | - | 47 |

**Table S3. Composition of telomeres and assembly gap in the *H. lupulus* cv. 10-12 (male) haploid and diploid genomes.**

| **Haploid** | Telomere | | |
|---|---|---|---|
| Chromosome | 5' end | 3' end | Gaps (count) |
| chr1 | Detect | Detect | 0 |
| chr2 | Detect | - | 0 |
| chr3 | Detect | Detect | 0 |
| chr4 | Detect | Detect | 0 |
| chr5 | Detect | Detect | 0 |
| chr6 | - | Detect | 0 |
| chr7 | Detect | - | 0 |
| chr8 | Detect | - | 0 |
| chr9 | Detect | Detect | 0 |
| chrX | - | Detect | 2 |
| chrY | - | Detect | 0 |

| **Diploid** | Telomere | | |
|---|---|---|---|
| Chromosome | 5' end | 3' end | Gaps (count) |
| chr1_hap1 | - | Detect | 4 |
| chr1_hap2 | Detect | - | 0 |
| chr2_hap1 | - | - | 21 |
| chr2_hap2 | Detect | - | 36 |
| chr3_hap1 | - | - | 7 |
| chr3_hap2 | - | - | 5 |
| chr4_hap1 | - | Detect | 7 |
| chr4_hap2 | - | - | 4 |
| chr5_hap1 | - | Detect | 19 |
| chr5_hap2 | - | - | 23 |
| chr6_hap1 | Detect | - | 2 |
| chr6_hap2 | - | - | 1 |
| chr7_hap1 | Detect | - | 6 |
| chr7_hap2 | Detect | Detect | 5 |
| chr8_hap1 | - | - | 1 |
| chr8_hap2 | - | - | 0 |
| chr9_hap1 | - | - | 9 |
| chr9_hap2 | - | - | 9 |
| chrX_hap1 | - | Detect | 52 |
| chrX_hap2 | Detect | - | 47 |

**Table S4. Composition of telomeres and assembly gap in the *H. japonicus* haploid genome.**

| Chromosome | Telomere | | Gaps (count) |
|---|---|---|---|
| | 5' end | 3' end | |
| chr1 | - | - | 2 |
| chr2 | - | Detect | 0 |
| chr4 | - | - | 1 |
| chr5 | - | Detect | 2 |
| chr6 | Detect | Detect | 1 |
| chr7 | Detect | - | 1 |
| chr8 | Detect | - | 0 |
| chrX | - | Detect | 15 |
| chrY1 | - | - | 26 |
| chrY2 | - | - | 30 |

**Table S5. Candidates of the sex-determining genes shared in the oldest strata of two *Humulus* species and *Cannabis sativa*. Expression levels in flowering buds and their bias between male and female were given.**

See excel sheet labeled with table S5

### Table S6. Phenotypic variations in p*EXER-EXER*-induced *N. tabacum*

| line | Phenotypic severity | Fillament length | pollen function | pistil length | petal length | flower numbers |
|---|---|---|---|---|---|---|
| pEXER-EXER-1 | - | no change | no change | no change | no change | no change |
| pEXER-EXER-2 | + | no change | less pollen | no change | no change | no change |
| pEXER-EXER-3 | +++ | very short | less pollen and no pollen function | longer | very short | less |
| pEXER-EXER-4 | ++ | short | less pollen and no pollen function | no change | short | less |
| pEXER-EXER-5 | + | no change | less pollen | no change | no change | no change |
| pEXER-EXER-6 | - | no change | no change | no change | no change | less |
| pEXER-EXER-7 | +++ | very short | less pollen and no pollen function | longer | very short | less |
| pEXER-EXER-8 | + | no change | less pollen | no change | no change | no change |
| pEXER-EXER-9 | ++ | short | less pollen and no pollen function | no change | no change | less |
| pEXER-EXER-10 | ++ | short | less pollen and no pollen function | no change | short | no change |
| pEXER-EXER-11 | - | no change | no change | no change | no change | less |
| pEXER-EXER-12 | - | no change | no change | no change | no change | no change |
| Cont.-1 | - | no change | no change | no change | no change | no change |
| Cont.-2 | - | no change | no change | no change | no change | no change |
| Cont.-3 | - | no change | no change | no change | no change | no change |
| Cont.-4 | - | no change | no change | no change | no change | no change |
| Cont.-5 | - | no change | no change | no change | no change | no change |
| Cont.-6 | - | no change | no change | no change | no change | no change |
| Cont.-7 | - | no change | no change | no change | no change | no change |
| Cont.-8 | - | no change | no change | no change | no change | no change |
| Cont.-9 | - | no change | no change | no change | no change | no change |
| Cont.-10 | - | no change | no change | no change | no change | no change |

**Table S7. List of differentially expressed genes in STS-treated female *H. japonicus*.**

See excel sheet labeled with table S7

**Table S8. Enriched GO terms in the genes down-regulated by STS treatment**

See excel sheet labeled with table S8

**Table S9. Phenotypic variations in male** H. japonicus **treated with ethephon or ACC**

| | Phnotypic severity | feminized inflorescence ratio[a] |
|---|---|---|
| **Ethephon-treatment to male plants** | | |
| Ethephon-1 | +++ | 0.889 |
| Ethephon-2 | +++ | 0.857 |
| Ethephon-3 | ++ | 0.444 |
| Ethephon-4 | +++ | 0.750 |
| Ethephon-5 | + | 0.200 |
| Cont.-1 | - | 0.000 |
| Cont.-2 | - | 0.000 |
| Cont.-3 | - | 0.000 |
| Cont.-4 | - | 0.000 |
| Cont.-5 | - | 0.000 |
| **ACC-treatment to male plants** | | |
| ACC-1 | + | 0.091 |
| ACC-2 | + | 0.125 |
| ACC-3 | - | 0.000 |
| ACC-4 | - | 0.000 |
| ACC-5 | ++ | 0.286 |
| Cont.-1 | - | 0.000 |
| Cont.-2 | - | 0.000 |
| Cont.-3 | - | 0.000 |
| Cont.-4 | - | 0.000 |
| Cont.-5 | - | 0.000 |

[a] Only top 1-3 flowers of spike inflorescence (flower nos > 10) exhibited perfect feminization.

### Table S10. Primer note

| Primer name | sequence (5'-3')[a] | |
|---|---|---|
| **For construction of *N. tabacum* transformation vector (*pHjEXER-HjEXER*)** | | |
| pPLV2-XEtRec-startF-BamH1 | **CCAACTCCATAAGGATC**ATGGCTCAATCATTCATCATGGAG | Target *BamH* 1 site in pPLV02 |
| pPLV2-XEtRec-stopR-BamH1 | **ACGATCGGGGATCGGATC**TCAATGCTCTAGTAATTCTGCAAGAACATTCC | |
| pPLV2-XEtRec-promF | **TTCTAGTTGGAATGGGTT**TTGAGTTCCCGAACCCTCTAGAA | Target *HpaI* site in pPLV02 |
| pPLV2-XEtRec-stopR | **TCCTTATGGAGTTGGGTT**TCAATGCTCTAGTAATTCTGCAAGAACATTCC | |
| **For construction of ALSV vector targeting *EXER* genes in *Humulus*** | | |
| R2m3hpRecB-IFF | GGGCCAGACCTCGAGACCATGCATTCAAGAACTGT | |
| R2m3hpRecB-IFR | TTGTCCCTCGGATCCGCCTGTCTCTTCTTGTGTG | |
| R2m3-ivF | GGATCCGAGGGACAAGGC | |
| R2m3-ivR | CTCGAGGTCTGGCCCCTG | |

[a] Bold underlined sequences are adapter sequences used in InFusion reaction.
